# Multi-location trials and population-based genotyping reveal high diversity and adaptation to breeding environments in a large collection of red clover

**DOI:** 10.1101/2022.12.19.520744

**Authors:** Michelle M. Nay, Christoph Grieder, Lea A. Frey, Helga Amdahl, Jasmina Radovic, Libor Jaluvka, Anna Palmé, Leif Skøt, Tom Ruttink, Roland Kölliker

## Abstract

Red clover (*Trifolium pratense* L.) is an outcrossing forage legume that has adapted to a wide range of climatic and growing conditions across Europe. Red clover is valued for its high yield potential and forage quality. The high amount of genetic diversity present in red clover provides an invaluable, but often poorly characterized resource to improve key traits such as yield, quality, and resistance to biotic and abiotic stresses. In this study, we examined the genetic and phenotypic diversity within a diverse set of 395 diploid red clover accessions via genome wide allele frequency fingerprinting and multi-location field trials across Europe. We found that the genetic structure of accessions mostly reflected their geographic origin and only few cases were detected, where breeders integrated foreign genetic resources into their local breeding pools. Phenotypic performance of accessions in the multi-location field trials revealed a very strong accession x location interaction. Notably, breeding material and cultivars generally performed well at the location where they were developed. Local adaptation was especially prominent in Nordic red clover accessions that showed a distinct adaptation to the growing conditions and cutting regime of the North. Our results confirmed that red clover cultivars were bred from regional ecotypes and show a narrow adaptation to regional conditions. Our study can serve as a valuable basis for identifying interesting material that express the desired characteristics and contribute to the adaptation of red clover to future climatic conditions.

## 1 Introduction

Red clover (*Trifolium pratense* L.) is the most important perennial forage legume in Northern and Central Europe. It is either grown in pure stands or in mixtures with tall grasses. Due to its ability to fix atmospheric nitrogen, red clover produces protein-rich forage and, when grown in mixtures, can provide additional nitrogen to companion grasses (Broderick, 1995; Taylor and Quesenberry, 1996; Halling et al., 2004; Nyfeler et al., 2011). Grass-legume mixtures have the additional benefit of reducing nitrogen leaching, increasing forage quality, and improving drought resilience (Lüscher et al., 2014; Hofer et al., 2016). Red clover is, therefore, an important component of sustainable grassland-based animal production and can substantially contribute to the protein self-sufficiency of Europe.

Red clover is native to Europe, the Middle East, North Africa and Central Asia, and was introduced to most temperate regions in the world as a forage crop (Taylor and Quesenberry, 1996; Boller et al., 2010). The benefits of red clover were readily appreciated by farmers, when they realized that the replacement of fallow by clover led to higher yields and better nutrient availability in the fields (Weir, 1926; Taylor and Quesenberry, 1996). Consequently, farmers started to harvest red clover seeds on farm for sowing in the crop rotation or for improving existing grasslands. This has led to the development of numerous landraces adapted to the specific management regimes and environmental conditions of the sites they emerged from (Taylor, 2008). Many of these landraces survived as naturalized populations, were integrated in breeders’ collections, or were conserved in genebanks. Together with other germplasm collections, these landraces laid the foundation for systematic breeding of red clover, which started in the second half of the 20^th^ century and is nowadays conducted in research organizations or small to large multinational companies throughout Europe.

Red clover is a naturally diploid (2n = 2x = 14), insect pollinated, outbreeding species with a high degree of self-incompatibility. Tetraploid red clover accessions have been produced through colchicine treatment, but despite their higher yield, their market success is hampered by low seed yield (Boller et al., 2010). In order to maintain a certain level of genetic diversity to maximize adaptability to a broad range of environmental conditions and due to its self-incompatibility, red clover is usually bred as population-based cultivars. Breeding schemes are mainly based on recurrent phenotypic selection either to directly develop open pollinated cultivars through population-improvement or to select suitable parental plants for mutual intermating (poly-crossing) and the development of synthetic cultivars (Posselt, 2010). The base populations used for breeding may consist of ecotype or wild populations, landraces, breeding material as well as commercial cultivars and are continuously complemented to ensure sufficient diversity in target traits. Specific breeding goals may differ between individual programs, but general breeding targets include high biomass and seed yield, persistence over the desired cultivation period and disease resistance (Abberton and Marshall, 2005; Taylor, 2008; Riday, 2010). In the northern regions of Europe, particular emphasis is put on improving freezing tolerance and resistance to clover rot, two factors strongly associated with persistence. Clover rot, caused by the ascomycete *Sclerotinia trifoliorum* Eriks., is favored by cooler temperatures, high humidity and long snow cover, and can lead to severe overwintering damage (Saharan and Mehta, 2008). Freezing tolerance is associated with winter survival and is particularly pronounced in wild populations from northern regions (Zanotto et al., 2021). In the warmer climates of Central Europe, persistence is an important breeding target with the specific aim to extend the period of cultivation for red clover cultivars over several growing seasons. Based on landraces developed through decades of on-farm seed production, a particular type of red clover (‘Mattenklee’) evolved and a number of highly persistent and high yielding cultivars were developed from this type (Kölliker et al., 2003; Boller et al., 2010). In the same region, resistance to southern anthracnose has gained importance as a breeding target. The disease is caused by *Colletotrichum trifolii* Bain & Essary and has benefitted from warmer summer temperatures, which has made it a limiting factor of red clover production in the warmer regions of Europe (Boller et al., 2010). The disease has long been recognized as a major threat in Southern USA, where targeted selection resulted in cultivars largely resistant to southern anthracnose (Taylor, 2008). North American red clover accessions are also often distinguished by their characteristic hairiness that is thought to aid in preventing leafhopper damage (Pieters and Hollowell, 1937). In order to account for different utilization requirements, different growth types have been developed. Early flowering clovers are capable of vigorous regrowth and remain generative throughout the growing period. Late flowering clover, termed ‘single-cut’ or ‘mammoth’ clover in the USA, remain vegetative after the first cut and are generally more persistent (Bird, 1948). Other types, such as the Swiss ‘Mattenklee’, combine the rapid regrowth of early flowering red clover with the persistence of the late flowering types (Boller et al., 2010). Thus, the various breeding activities combined with the use of landraces and the introgression of wild populations for specific characteristics resulted in very diverse genetic resources, adapted to vastly different environments – ranging from arctic to mediterranean and continental climates. However, most of these breeding activities were focused on creating cultivars optimized for local or regional needs, and little effort was spent on breeding for a broader range of environments. The latter may be particularly important in the future in view of rapidly changing environmental conditions through climate change and shifts in priorities for land-use.

Plant genetic resources, *i.e*., the genetic material contained in wild relatives, ecotypes, landraces, and cultivars, are crucial for the continued improvement of modern forage crop cultivars (Boller and Greene, 2010). For red clover, the European Search Catalogue for Plant Genetic Resources (EURISCO; https://eurisco.ipk-gatersleben.de/apex/eurisco_ws/r/eurisco/home) currently lists more than 10,600 red clover accessions and many more are likely to be found in non-European or local germplasm collections. For targeted utilization in breeding programs, detailed phenotypic and genotypic characterization data is indispensable, but is often not available for many of the current genetic resources. Molecular genetics or DNA-based markers allow a rapid characterization of genetic diversity in a large number of samples, independent of environmental effects (Bachmann, 1994). PCR-amplification based marker systems targeting either many unspecific loci (*e.g*., amplified fragment length polymorphism (AFLP) markers; Vos et al., 1995) or single highly variable loci (*e.g*., simple sequence repeat (SSR) markers; Tautz, 1989) have long been the methods of choice for analyzing genetic diversity in forage crop species where genome sequence information has been scarce. More recently, highly cost-effective genotyping methods based on high-throughput sequencing of restriction site-associated DNA such as RAD-seq (Baird et al., 2008) or genotyping-by-sequencing (GBS; Elshire et al., 2011) have become available. These methods allow an even larger number of samples to be efficiently genotyped at several thousand loci. In red clover, molecular markers have been successfully used to investigate the origin of red clover types and cultivars (Semerikov et al., 2002; Kölliker et al., 2003), the genetic structure of germplasm collections (Herrmann et al., 2005; Dias et al., 2008; Gupta et al., 2017) and the distribution of genetic diversity over a broad geographic range (Jones et al., 2020). As expected for an outbreeding species, a large proportion of the genetic variation is observed between individuals within populations (Kölliker et al., 2003; Dias et al., 2008), making it necessary to analyze multiple individuals to capture the genetic diversity present in a population. Pooling leaves of 20 individual red clover plants prior to DNA extraction has been proven effective to analyze genetic diversity on a population level (Herrmann et al., 2005). In perennial ryegrass, GBS on pooled leaf samples has proven useful to establish genome wide allele frequency fingerprints and to differentiate populations (Byrne et al., 2013). Despite the large number of accessions available in genebanks, most studies have used a rather restricted number of accessions and/or individual plants or focused on specific geographic regions. Jones et al. (2020) investigated genetic variability of 75 red clover accessions using eight to 16 individual plants per accession and more than 8,000 single nucleotide polymorphism (SNP) markers. The accessions were clearly grouped into four groups, corresponding to their geographic origin (Asia, Iberia, UK and Central Europe). The largest study so far investigated genetic diversity of 382 Nordic red clover accessions using 661 SNPs (Osterman et al., 2022).

Although studies on genetic diversity can substantially assist breeding decisions, knowledge about phenotypic diversity is key to efficiently exploit red clover germplasm for cultivar development. While some studies have investigated the variability and inheritance of specific traits such as disease resistance (Frey et al., 2022), seed yield (Herrmann et al., 2005) or flowering time (Jones et al., 2020), only a few studies have investigated phenotypic diversity of several traits within a region or country and even fewer have studied diversity across regions. A study by Zanotto et al. (2021) that was conducted with 48 ecotypes and six cultivars from Norway, Sweden, and Finland found large variation in winter survival and yield between the accessions. Wild accessions sometimes outperformed commercial cultivars, indicating their value for improving adaptation to colder climates. Winter survival was found to vary considerably between the four locations and could only partially be predicted by a test under controlled conditions (Zanotto et al., 2021). Furthermore, in a meta-analysis of legume yields in trials conducted in several locations ranging from Southern Germany to Northern Norway, Halling et al. (2004) found large differences in forage yield not only across different environments but also between early, intermediate and late flowering accessions. These findings highlight the need to account for accession x location interactions and to characterize breeding germplasm in their respective environments.

In order to provide a solid foundation for the utilization of red clover genetic resources in breeding programs, we aimed at (i) establishing a representative, diverse collection of red clover accessions, (ii) characterizing the genetic diversity and structure present in this collection, (iii) assessing the phenotypic diversity of agriculturally relevant traits at multiple locations and (iv) getting an insight into the extent of accession x location interaction for future adaptive breeding efforts.

## 2 Materials and Methods

### 2.1 Red clover diversity panel

In this study, the red clover diversity panel for the project ‘Breeding forage and grain legumes to increase EU’s and China’s protein self-sufficiency – EUCLEG’(www.eucleg.eu) was compiled to cover as much of the available genetic diversity present in red clover in Europe as possible. To do so, all members of the project consortium, including breeders, research institutes and genebanks, were asked to contribute with their available material. In addition, other institutes outside the consortium were contacted for possible contributions to compile as many accessions as possible. Accessions were delivered with specific material transfer agreements from each supplier to a central coordinator, who further distributed seeds under the same agreements to the researchers performing the experiments. The final panel consists of 395 accessions representing available diversity of diploid red clover in Europe, complemented with a few accessions from the Americas and Oceania (Figure 1, Supplementary Table S1). In addition to country, accessions were grouped by their region of origin: Americas (16 accessions), Oceania (10), Central Europe (51). Eastern Europe (82), Switzerland (81), Southern Europe (13), Northern Europe (139) and others (3, Japan, Iran).

**Figure 1:**
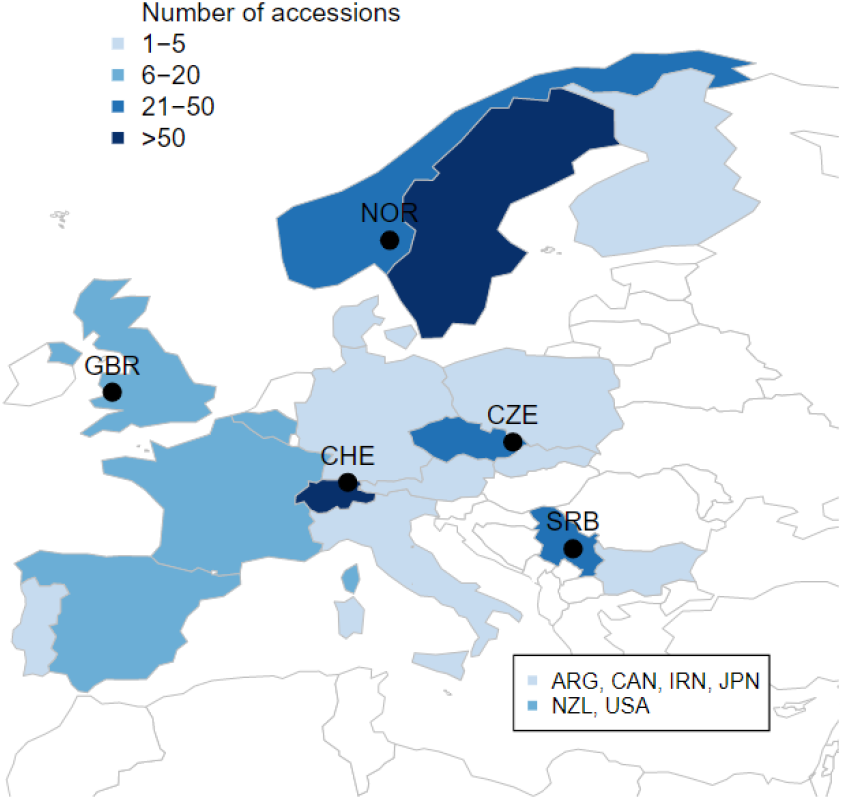
Origin and number of the European accessions in the EUCLEG red clover panel. Additional accessions from Argentina (ARG, 5), Canada (CAN, 1), Iran (IRN, 1), Japan (JPN, 2), New Zealand (NZL, 10) and USA (10) are not shown on the map. Locations of the field trials in Norway (NOR), Great Britain (GBR), Switzerland (CHE), Czech Republic (CZE) and Serbia (SRB) are indicated by black circles.

### 2.2 Genotyping

Genotyping of the 395 accessions and SNP calling was conducted as described in Frey et al. (2022). In brief, plants from all accessions were grown in the greenhouse in 96-compartment trays. Fresh leaves from 200 plants per accession were harvested at the one-leaf stage and pooled for DNA extraction. DNA extraction was done with the QIAGEN DNeasy 96 Plant kit (QIAGEN, Manchester, UK) according to the user manual. After normalizing to 20 ng μl^-1^, DNA of each accession pool was shipped to LGC Genomics (Berlin, Germany) for pooled-GBS and PE-150 Illumina sequencing. SNP calling and calculations of allele frequencies were done as described in Keep et al. (2020). A detailed description of the parameters used for this study are given in the supplementary methods of Frey et al. (2022). SNPs were only retained if allele frequencies of at least 10 accessions were between 0.05 and 0.95 and if mean allele frequencies across all accessions were between 0.05 and 0.95. SNP positions with more than 5% missing values were discarded. Remaining missing values in the allele frequency matrix were replaced by the mean allele frequency across all accessions at the given SNP position.

Genetic diversity among accessions was investigated using principal component analysis implemented in the function ‘prcomp’ (R Core Team, 2020). The genetic structure was further studied using discriminant analysis of principal components implemented in the ‘adegenet’ package version 2.7.3 (Jombart, 2008). The potential number of subpopulations (clusters) was estimated using successive K-means clustering. Partition of variance was investigated using permutational analysis of variance implemented in the function ‘adonis2’ of the ‘vegan’ package (Oksanen et al., 2022).

### 2.3 Field trials

#### 2.3.1 Locations and trial designs

Field plot trials with the full set of all 395 accessions were conducted at Agroscope in Tänikon, Switzerland (CHE 47.480°N, 8.904°E, 535 m.a.s.l.) and at DLF Seeds AS in Hladké Životice, Czech Republic (CZE 49.690°N, 17.960°E, 220 m.a.s.l.). A reduced set of 110 accessions was grown at Graminor in Bjørke, Norway (NOR 60.757°N, 11.203°E, 147 m.a.s.l.) and reduced sets of 100 accessions each were grown at Aberystwyth University in Aberystwyth, Wales (GBR 52.427°N, 4.020°W, 35 m.a.s.l.) and at the Institute for Forage Crops Kruševac Ltd. in Kruševac, Serbia (SRB 43.583°N, 21.206°E, 150 m.a.s.l.). To assess flowering time independently of the cutting regime of the main plot, additional observation rows with two replicates were sown at each trial location.

For the main plot trials in CZE, CHE and GBR, partially replicated (Prep) designs were employed, with accessions being grown unreplicated, in two or in six replicates (Table 1). For these trials, no complete blocks (CB) were applied, and each row and column constituted an incomplete block (IB1 and IB2, respectively). The main plot trials in NOR and SRB were designed as alpha designs with two CB containing the full set of accessions (*i.e*. complete replicates) and incomplete blocks in only one direction (IB1) of dimension 1 row by 10 columns (10 plots) in NOR and dimension 1 row by 5 columns (5 plots) in SRB. Observation rows for assessing flowering time were designed with two CB in all locations. Due to their larger size, the observation rows in CHE and CZE further included rows as first dimension of incomplete blocks (IB1) and columns as second dimension of incomplete blocks (IB2).

Due to different growing conditions and standard equipment used by the experimenters in the five locations, trials were set up and nursed according to local standard procedures. Plot size ranged between 5 and 8 m^2^ with a sowing density of 600 to 800 germinating seeds m^-2^ (Table 1). All trials were to be sown at the beginning of the 2018 field season to establish plant stands with minimal weed infestation for subsequent measurements during 2019 and 2020. Due to unsuitable weather conditions, sowing had to be postponed to 8^th^ of August in GBR and the trial in SRB had to be re-sown on the 17^th^ of September due to drought and flood destroying the first trial sown in spring. Because of additional damage to the field trial in SRB during early 2019, data could only be assessed from the first replicate at this location.

**Table 1.** Layout of field trials at the different locations.

| Parameter | Trial location |  |  |  |  |
| --- | --- | --- | --- | --- | --- |
|  | CHE | CZE | GBR | NOR | SRB |
| Design | Prep | Prep | Prep | Alpha | Alpha |
| Number of accessions |  |  |  |  |  |
| - with 1 plot | 350 | 350 | 80 | - | - |
| - with 2 plots | 35 | 35 | 15 | 109 | 100 |
| - with 6 or more plots | 10 | 10 | 5 | 1<sup>1)</sup> | - |
| - in total | 395 | 395 | 100 | 110 | 100 |
| Number of plots | 490<sup>2)</sup> | 480 | 140 | 240 | 200 |
| Trial dimension (rows x columns) | 7 x 70 | 20 x 24 | 14 x 10 | 12 x 20 | 20 x 10 |
| Complete blocks | No | No | No | Yes | Yes<sup>3)</sup> |
| Incomplete blocks 1<sup>st</sup> dimension | IB1 (1 x 70) | IB1 (1 x 24) | IB1 (1 x 10) | IB1 (1 x 10) | IB1 (1 x 5) |
| Incomplete blocks 2<sup>nd</sup> dimension | IB2 (7 x 1) | IB2 (20 x 1) | IB2 (14 x 1) | - | - |
| Plot size | 1.5 x 5m | 1.6 x 5m | 1.2 x 5m | 1.25 x 4m | 1 x 5m |
| Sowing density (seeds m<sup>-2</sup>) | 600 | 800 | 600 | 600 | 600 |
| Sowing date | 2018-04-21 | 2018-05-30 | 2018-08-08 | 2018-06-27 | 2018-09-17 |

1) Accession 'Lea' was grown with 22 plots in total (*i.e.* 11 plots per complete block)
2). 10 plots of the trial in CHE were sown with a control accession to facilitate harvest. No data were recorded for these plots (missing values).
3) No data assessment was possible in second complete block.

#### 2.3.2 Phenotyping

All traits were assessed on a plot basis during the year of sowing (Y0), and the first and second main harvesting years (Y1 and Y2, respectively). The cutting regime followed common practice for the respective locations, ranging from 3 (NOR), over 4 (CZE, GBR, SRB), to 5 (CHE) cuts per main harvesting year. Before each cut, different traits were assessed visually: plant vigor (VIG) was rated on a scale from 1 (very weak) to 9 (very vigorous), weed occurrence was rated as the percentage of plot biomass derived from weeds (Weed%) and the resistance to spontaneously occurring diseases was rated on a scale from 1 (very high disease infestation) to 9 (no disease infestation). Diseases observed in the trials were powdery mildew caused by *Microsphaera trifolii* (Grev.) U. Braun, southern anthracnose caused by *Colletotrichum trifolii* Bain & Essary, clover rot caused by *Sclerotinia trifoliorum* Eriks., plant rot caused by different *Fusarium ssp*. and brown spot caused by *Stemphylium sarciniforme* (Cav.) Wiltsh.

For each cut, plots were harvested with a plot harvester and weight of fresh matter was determined using a machine mounted balance. Fresh matter yield (FMY) was determined as fresh matter per plot divided by the plot area. In all trials but CHE, a subsample per plot was dried in an oven at 100°C to constant weight and percentage of dry matter concentration (DMC) was determined as weight of dry subsample divided by weight of fresh subsample multiplied by 100. For the trial in CHE, DMC was determined indirectly using near infrared spectroscopy (NIRS) measurements taken on the plot harvester. Dry matter yield (DMY) corrected for weed occurrence was then calculated for each plot as

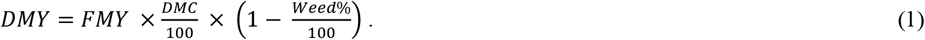

For the first and second cut of Y1, samples for forage quality analysis were taken from each plot at all locations and were dried to constant weight at 50 to 60 °C. Dried samples were then shipped to Agroscope for central processing. Grinding of samples was done using a cutting mill (SM200, Retsch, Haan, Germany) with a mesh size of 0.75 mm. NIR spectra on the ground samples were measured in the wavelength range from 800 to 2500 nm using a laboratory spectrometer (NIRFlex N-500, Büchi, Flawil, Switzerland). The two parameters crude protein content (CP) and digestible organic matter (DOM) were then determined for each sample using NIR-calibrations developed in-house based on reference values determined in red clover with the respective reference methods. Reference measurements for calibration development of CP (in % of dry matter) were established by multiplying total nitrogen (% of dry matter) determined following Kjeldahl (1883) by a factor of 6.25. Reference determination for DOM (in mg g^-1^) was performed using the *in vitro* assay following Tilley and Terry (1963).

In addition to the traits assessed before each cut, plant density was determined visually at the end of Y0 and at the beginning and end of Y1 and Y2. At each date, plant density was rated in percentage of the most dense plot observed at the end of Y0. Date of flowering (DOF) was determined in Y1 and Y2 as the number of days after 1^st^ of January when at least five plants per plot started flowering (visible appearance of pink color of the petals). Because the plots were cut before all accessions would start flowering, DOF was determined on separate observation rows. In CHE, DOF was assessed in Y2, whereas it was assessed in Y1 for all other locations. For accessions that did not start flowering at all, a value of 222 was assigned.

#### 2.3.3 Statistical Analyses

Traits were analyzed for each cut separately. For comparison among locations, additional secondary traits were calculated. Mean vigor rating for Y1 and Y2 (VIG_Y1 and VIG_Y2, respectively) was calculated as simple mean over all cuts and a missing value was assigned if data from one cut was not available. The sum of DMY for Y1 and Y2 (DMY_Y1 and DMY_Y2, respectively) was calculated as sum of DMY over all cuts of the respective year.

Locations were analyzed separately using the following model:

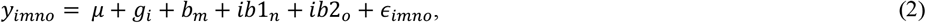

where *y_immo_* represents the observation for trait *y* on a single plot basis, *μ* denotes the overall mean, *g_i_* is the effect of accession *i*, *b_m_* the effect of block *m*, *ib*1_*n*_ the effect of incomplete block 1 (*i.e*. row) *n*, and *ib*2_*o*_ the effect of incomplete block 2 (*i.e*. column) *o*, and *ε_imno_* is the residual error. If not present in the design of a given location, the respective factor was omitted for analysis (i.e. *b_m_* for CHE, CZE and GBR and *ib*2_*o*_ for NOR). Because data was only available from one complete block (*i.e*. replicate) in SRB, no separate analysis could be performed for this location. In a first model, to estimate the best linear unbiased estimates of accession means (BLUEs), *ib*1_*n*_ and *ib*2_*o*_ were considered as random, while all other effects were considered as fixed. In a second model, to estimate the best linear unbiased predictors of accession means (BLUPs) and the respective variance components, *g_i_* was additionally considered as random. For all subsequent phenotypic analyses, BLUE values were used.

Multi-location analyses using data from all five locations were done following the model:

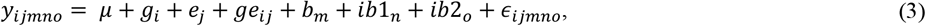

where *y_ijmno_* represents the observation for trait *y* on a single plot basis, *μ* denotes the overall mean, *g_i_* is the effect of accession *i*, *e_j_* the effect of the location *j*, *ge_ij_* the interaction between accession *i* and location *j*, *b_m_* the effect of complete block *m* (if available), *ib*1_*n*_ the effect incomplete block 1 *n*, *ib*2_*o*_ the effect of incomplete block 2 *o*, and *ε_ijmno_* is the residual error. In a first model, effect *ib*1_*n*_ and *ib*2_*o*_ were considered as random, while all other effects were considered as fixed to estimate BLUEs. To estimate respective variance components, *g_i_* and *ge_ij_* were additionally considered as random in a second model. For single and across location analyses, heritability (h^2^) values were calculated according to the method of Walsh and Lynch (2018) as the slope from the linear regression of BLUPs on BLUEs of a given trait.

For the principal component analysis (PCA) on phenotypic data, BLUEs of the traits DOF_Y1, CP_Y1.C1, CP_Y1.C2, DMY_Y1.C1, DMY_Y1.C2, DMY_Y1, DMY_Y2, VIG_Y1, VIG_Y2, PD_Y1 were used. Diseases were additionally included, if they occurred at a given location. Occurring diseases were southern anthracnose and powdery mildew in CHE, brown spot in SRB, fusarium in CZE and clover rot in NOR. Only accessions with data in at least 50% of the selected traits were included and only traits that have data in at least 80% of the accessions were kept. Hence, DOF, DMY_Y2 and VIG_Y2 were not included for NOR and DMY_Y2 and VIG_Y2 were not included for CZE in the analysis. The remaining missing values were imputed with the mean of the trait.

For estimation of variance components and calculation of mean values, heritability, correlations coefficients, and PCA, DOF values of accessions that did not flower were set to ‘not available’. Thereby, an artificial inflation of variance could be omitted. For all graphical representations, the preset value of 222 was kept for these accessions.

All statistical analyses were conducted with R version 4.1.2 within RStudio v4.0.5. (R Core Team, 2020; RStudio Team, 2021) using the functions ‘lmer’ and ‘ranef of package ‘lme4’ and ‘emmeans’ of package ‘emmeans’ for mixed model analyses (Bates et al., 2015). For PCA analysis, the function ‘prcomp’ of package ‘stats’ was used. In addition, the packages ‘tidyverse’, ‘psych’, ‘ggfortify’, ‘rworldmap’, ‘gridExtra’, ‘cowplot’, ‘grid’ and ‘ggpubr’ were used for data handling and illustration.

## 3 Results

### 3.1 Genetic diversity of the EUCLEG red clover panel

Genotyping-by-sequencing on pooled samples of the 395 accessions resulted in a set of 20,137 quality controlled and filtered SNPs evenly distributed across the red clover genome (Frey et al., 2022). Three samples (TP107, TP133, TP309; Supplementary Table S1) did not meet the quality requirements and were excluded from further analyses, resulting in a reduced dataset of 392 accessions for which GBS data was available.

The first two axes of the PCA together explained 14.5% of the genetic variation and allowed to distinguish some of the accessions based on their region of origin and their type (Figure 2). Principal component 1 (PC1) mainly separated ecotypes from Southern Europe from the remaining accessions, while PC2 allowed to distinguish accessions from Switzerland, Central Europe and Northern Europe. This separation was even more pronounced for PC3, which also allowed to distinguish breeding material and cultivars from the ecotypes and landraces in the Northern European material (data not shown). Permutational multivariate analysis of variance identified 16.8% of the variation to differentiate among regions, while 12.4% was attributed to variation between countries of origin and 70.8% to variation among accessions within countries (Table 2).

**Figure 2:**
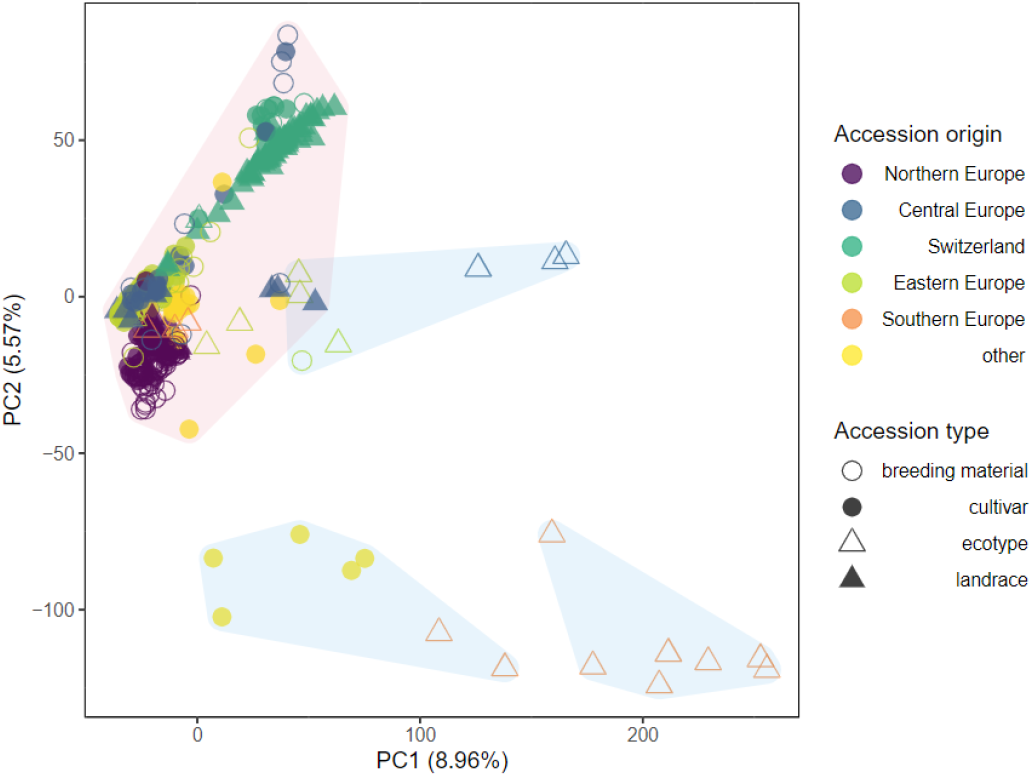
Genetic similarity of 392 red clover accessions within the EUCLEG panel revealed by principal component analysis of allele frequency data based on 20,137 SNP markers and pooled leaf samples of 200 individuals per accession. Colors indicate the region of origin of the accessions and symbols indicate the accession type. K-means clustering identified three major clusters, which combined contained 371 accessions (shaded in red) and three minor clusters each comprising seven accessions (shaded in blue).

**Table 2:** Permutational multivariate analysis of variance of 392 red clover accessions from 23 countries and six regions genotyped using 20,137 SNPs.

| Source of variation | DF<sup>1</sup> | SS<sup>2</sup> | Variance explained (%) | *F*-value | *P*-value |
| --- | --- | --- | --- | --- | --- |
| Region | 5 | 16,423 | 0.170 | 17.694 | 0.091 |
| Origin (region) | 17 | 11,956 | 0.123 | 3.789 | 0.091 |
| Accessions (origin) | 369 | 68,498 | 0.707 |  |  |

<sup>1</sup> Degrees of freedom; <sup>2</sup> Sum of squares

Successive K-means clustering of the PCs and comparing the models using the Bayesian Information Criteria (BIC) indicated the existence of six to 11 clusters (Supplementary Figure S1). Defining the clusters using K = 6 (the number of regions in the dataset) revealed three minor clusters that together contained 21 accessions, while the remaining three clusters contained the remaining 371 accessions (Figure 2). Consequently, a second PCA was performed on the subset of the 371 accessions grouped in the three major clusters.

PCA of the 371 accessions grouped in the three main clusters by successive K-means clustering further separated accessions according to their origin and accession type (Figure 3). PC1 and PC2 allowed to clearly separate three major groups: accessions from Northern Europe, accessions from Switzerland and accessions from the remaining regions. Particularly, PC2 distinguished Northern European breeding material and cultivars from landraces, while for Swiss accessions, cultivars are separated from landraces (Figure 3).

**Figure 3:**
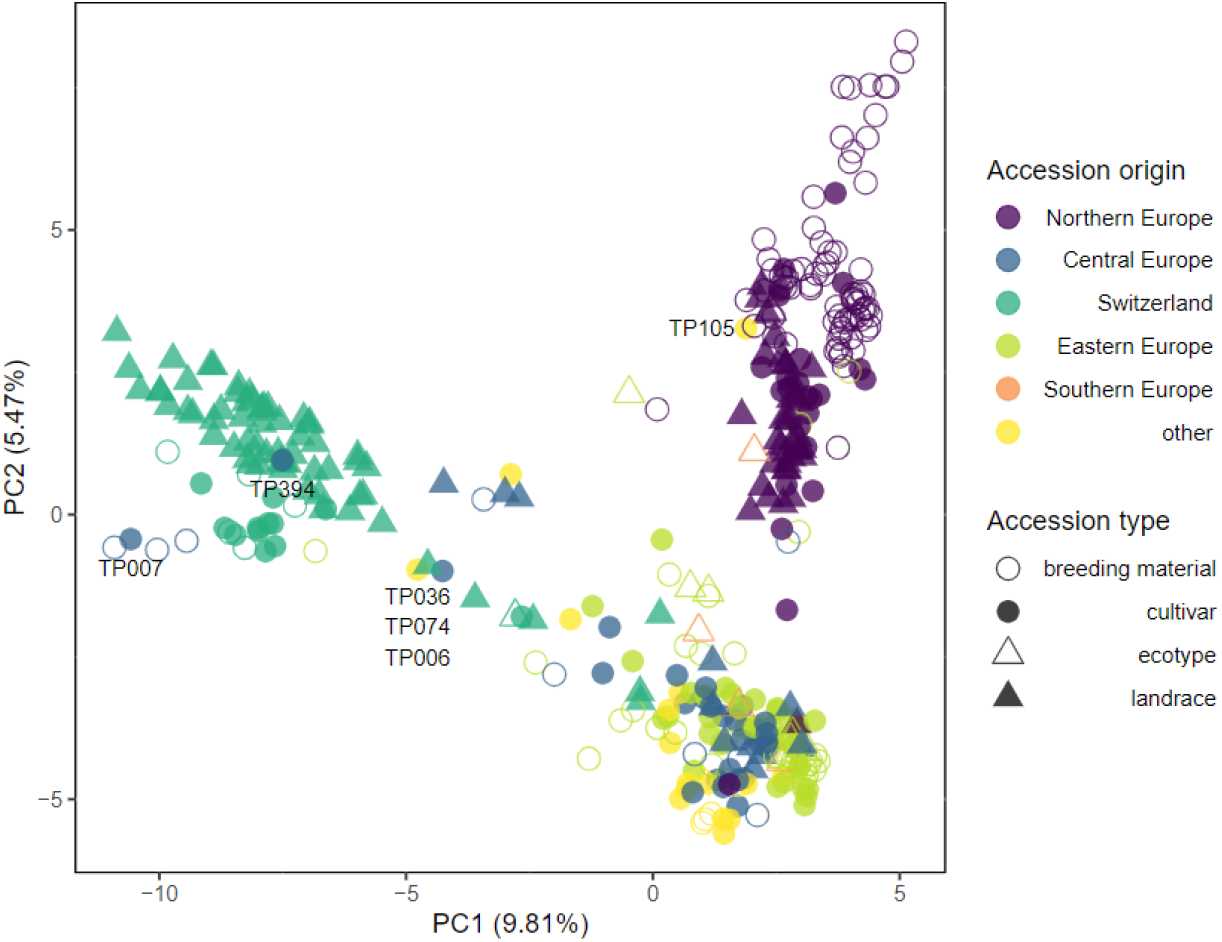
Genetic similarity of 371 selected red clover accessions revealed by principal component analysis of allele frequency data based on 20,137 SNP markers and pooled leaf samples of 200 individuals per accession. Accessions forming the three major clusters of the complete set identified by K-means clustering (Fig. 2) were selected. Colors indicate the region of origin of the accessions and symbols indicate the accession type. Accession IDs are given for selected accessions.

### 3.2 Field trials

With the exception of the second field replicate in SRB, all trials were successfully established during the initial year (Y0) and could be used for assessment of yield, quality and disease occurrence data in the subsequent main harvesting years (Supplementary Table S2). In CHE, DMY was highest during the first three cuts in Y1 and strongly decreased thereafter (Figure 4). Due to southern anthracnose infections during cut 4, plant density was reduced allowing weeds to spread. In Y2, DMY in CHE started at a lower level compared to Y1 and was reduced to nearly zero for most plots towards the end of the season with red clover plants being replaced by weeds. In CZE, DMY for the first cut of Y1 (DMY_Y1.C1) was highest among all locations, but strongly decreased with the subsequent cut (Figure 4, Table 3). Along with the observation of severe fusarium rot during the third cut and subsequent infestation with mice, no more yield assessment was possible in CZE and the trial had to be abandoned. In GBR, weed infestation started at a relatively high level at cut 1 in Y1, but continuously decreased until cut 3, staying relatively low for the remaining time of the experiment. DMY thereby increased, reaching highest values for cut 3 in Y1 and for cut 2 in Y2. In NOR, three cuts were performed in both Y1 and Y2 with the third cut having a reduced DMY compared to the first two cuts in both main harvesting years. Occurrence of clover rot was observed before the first cut in Y1. Weed occurrence increased after winter at cut 1 in Y2, but again decreased thereafter. In SRB, plots had to be resown in autumn of Y0 and were cut four times in Y1, with DMY values decreasing with subsequent cuts. In Y2, DMY started at a higher level compared to Y1 and also decreased until the third and last cut. Due to a generally low infestation with weeds, no weed infestation was assessed in SRB.

**Figure 4:**
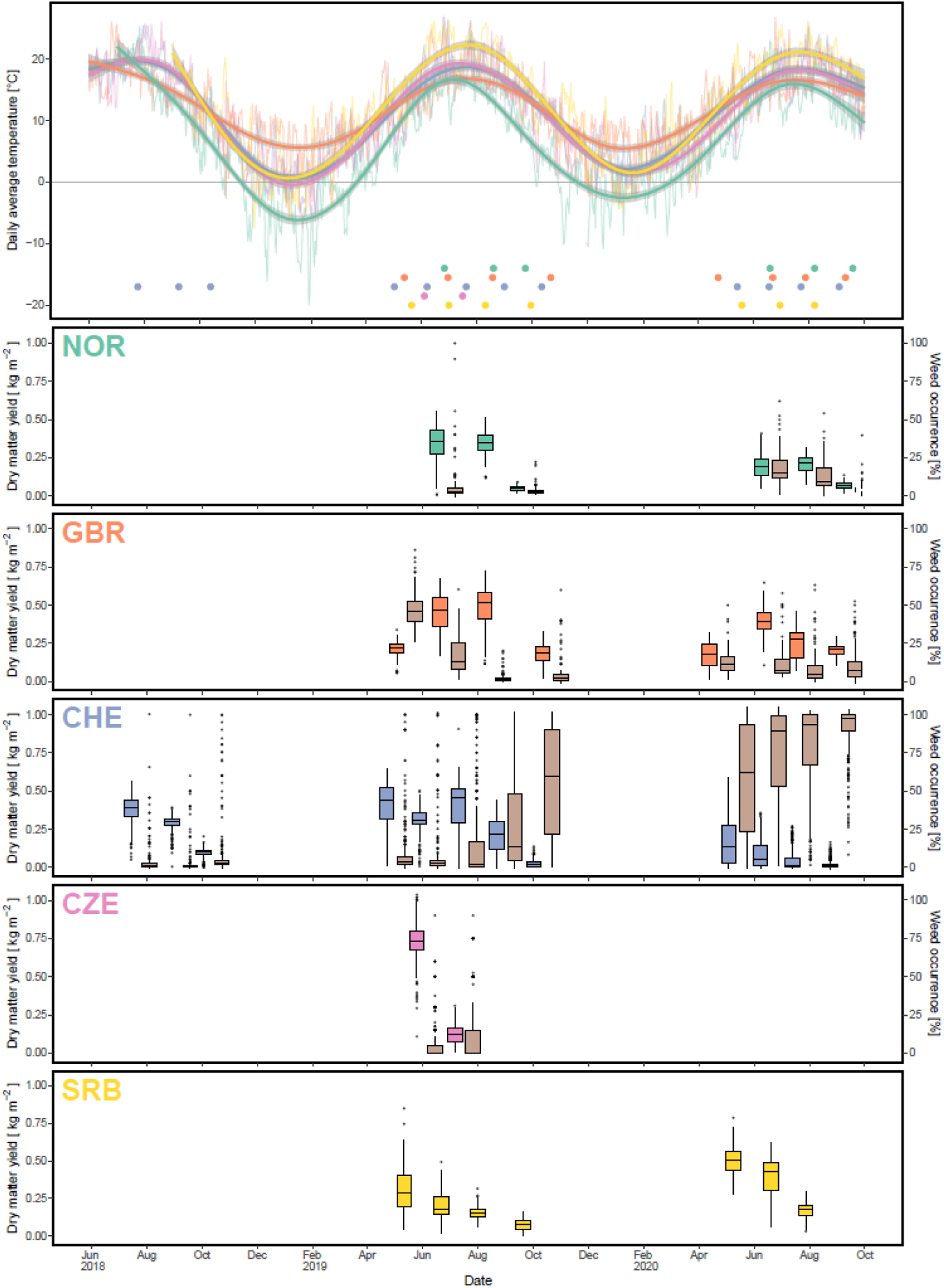
Uppermost panel: temperature course during the period of the experiment with thin lines denoting the daily average temperature, thick lines denoting the smoothed temperature and dots showing the cutting dates per location. Subsequent panels: boxplots of dry matter yield (different colour per location) and weed occurrence (brown colour) for each cut and location.

Due to the lack of data from location CZE after cut 2 in Y1 and the heavily reduced plant stands at location CHE after cut 4 in Y1 due to non-adapted (*i.e*., southern anthracnose susceptible) accessions coming along with a high weed occurrence, subsequent analyses mostly focused on Y1. Given the vast amount of data obtained, a main focus was on the traits date of flowering in Y1 (DOF_Y1), total dry matter yield in Y1 and Y2 (DMY_Y1 and DMY_Y2, respectively), average vigor of Y1 (VIG_Y1) and plant density measured in autumn of Y1 (PD_Y1). In addition, we took a closer look at the crude protein data of the first and second cut in Y1 (CP_Y1.C1, CP_Y2.C1). Furthermore, the natural occurrence of diseases in the different trials is reported and discussed

**Table 3.** Mean and range of best linear unbiased estimators (BLUE) per accession observed at the different locations. Traits are dry matter yield (DMY), crude protein content (CP), digestible organic matter (DOM), date of flowering (DOF), plant density (PD) and plant vigor (VIG). Extensions _Y1 and _Y2 denote the first and second year of observation, .C1 and .C2 the first and second cut per year of observation, respectively.

| Parameter | NOR |  | GBR |  | CHE |  | CZE |  | SRB |  |
| --- | --- | --- | --- | --- | --- | --- | --- | --- | --- | --- |
|  | Mean | Range | Mean | Range | Mean | Range | Mean | Range | Mean | Range |
| DMY_Y1 (kg m<sup>-2</sup>) | 0.74 | 0.19-1.01 | 1.33 | 0.44-1.83 | 1.34 | 0.01-2.03 | 0.86 | 0.12-1.26 | 0.74 | 0.14-1.44 |
| DMY_Y1.C1 (kg m<sup>-2</sup>) | 0.34 | 0-0.55 | 0.21 | 0.05-0.34 | 0.41 | 0.01-0.65 | 0.73 | 0.11-1.07 | 0.31 | 0.05-0.85 |
| DMY_Y1.C2 (kg m<sup>-2</sup>) | 0.34 | 0.12-0.51 | 0.45 | 0.17-0.67 | 0.31 | (-0.03)-0.5 | 0.12 | (-0.02)-0.31 | 0.20 | 0.02-0.49 |
| DMY_Y2 (kg m<sup>-2</sup>) | 0.50 | 0.18-0.85 | 1.03 | 0.41-1.42 | 0.28 | (-0.06)-1.24 | NA | NA | 1.07 | 0.4-1.55 |
| DMY_Y2.C1 (kg m<sup>-2</sup>) | 0.20 | 0.05-0.41 | 0.17 | 0.01-0.32 | 0.15 | (-0.04)-0.58 | NA | NA | 0.50 | 0.28-0.79 |
| DMY_Y2.C2 (kg m<sup>-2</sup>) | 0.21 | 0.07-0.31 | 0.40 | 0.11-0.65 | 0.07 | (-0.02)-0.36 | NA | NA | 0.38 | 0.06-0.62 |
| CP_Y1.C1 (g kg<sup>-1</sup>) | 174 | 119.73-215 | 208 | 147-265 | 249 | 194-300 | 162 | 93-250 | 151 | 105-225 |
| CP_Y1.C2 (g kg<sup>-1</sup>) | 172 | 120-226 | 173 | 116-220 | 247 | 179-295 | 200 | 122-266 | 153 | 91-204 |
| DOM_Y1.C1 (mg g<sup>-1</sup>) | 584 | 478-679 | 731 | 661-800 | 682 | 584-738 | 624 | 545-698 | 525 | 367-622 |
| DOM_Y1.C2 (mg g<sup>-1</sup>) | 576 | 525-621 | 647 | 532-731 | 638 | 523-748 | 524 | 444-603 | 531 | 375-630 |
| DOF_Y1 (d) | 181 | 167-199 | 150.8 | 109-222 | 153.9 | 121-222 | 153.2 | 142-164 | 132.4 | 125-142 |
| PD_Y1.beginning (%) | 26.1 | (-0.1)-66.2 | 51.5 | 4.8-96.7 | NA | NA | 7.54 | 1.25-9.53 | 97.6 | 90-100 |
| PD_Y1.end (%) | 53.4 | 5.08-84.7 | 57.8 | 17.9-74.4 | 41.9 | (-7.7)-94.9 | 3.42 | (-0.5)-9.62 | 91.8 | 80-100 |
| VIG_Y1 (rating 1-9) | 8.36 | 6.55-9.16 | 6.53 | 3.25-8.01 | 5.63 | 2.02-8.46 | 6.11 | 1.8-8.01 | 5.74 | 2.25-8.75 |
| VIG_Y2 (rating 1-9) | 7.91 | 6.85-8.78 | 6.96 | 4.48-8.74 | 2.66 | 0.69-8.12 | NA | NA | 5.67 | 3.4-7.2 |

#### 3.2.1 Variance components and heritability

Quality parameters of the red clover herbage were measured for the first and second cut of Y1. From single location analysis, most of the variation in CP was explained by the residual variance and only minor effects of accession or field heterogeneity could be observed (Figure 5, Supplementary Table S2). The second quality parameter, DOM, showed a very similar pattern to CP and was also largely dominated by residual variance (Supplementary Table S2). In comparison, variation in DMY of the corresponding cuts (DMY_Y1.C1, DMY_Y1.C2) could, except for location CZE, be explained to a large part by accession. Also, for DOF_Y1, PD_Y1, DMY_Y1, DMY_Y2, and VIG_Y1, a large proportion of the observed variation could be attributed to the accession, while the variance, due to field variation, indicated by block, and the residual variance were relatively low (Figure 5). When all locations were considered simultaneously in a multi-location analysis, a large accession x location effect was observed for most traits. For CP, values observed for the first cut (CP_Y1.C1) were still dominated by residual variance, while the accession variance observed for the second cut (CP_Y1.C2) was replaced by accession x location interaction variance. For DMY at specific cuts, *e.g*., DMY_Y1.C1 and DMY_Y1.C2, the accession variance observed at single locations was almost entirely replaced by accession x location interaction variance. For sum of DMY per main harvest year, *e.g*., DMY_Y1 and DMY_Y2, and average vigor of year 1 (VIG_Y1) some variance attributed to accessions was still observed in the multi-location analysis. The absolute amount of variation varied considerably between trials and traits. For the trial in SRB, no variance components could be estimated, because data could only be obtained from one replicate per accession.

**Figure 5:**
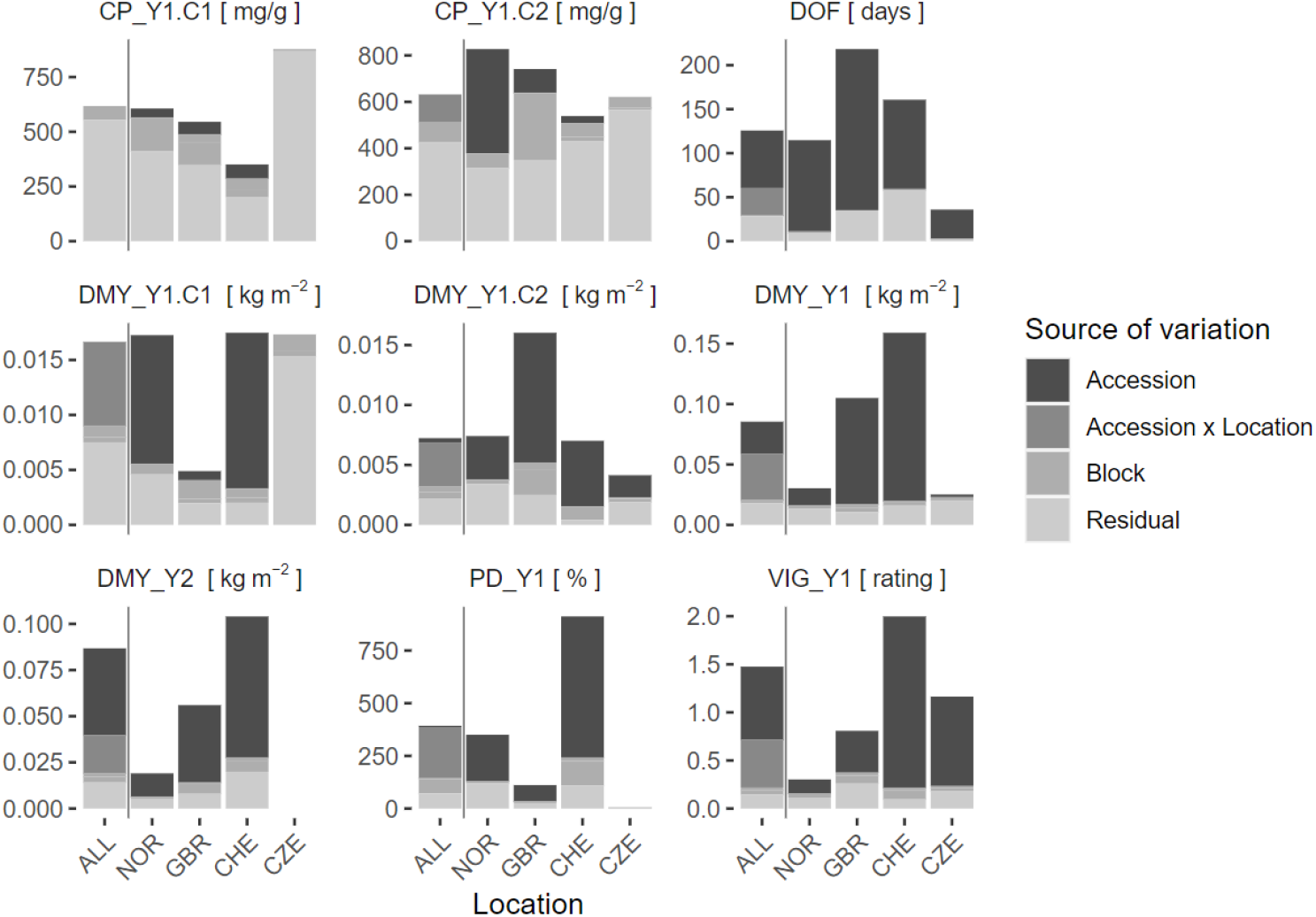
Variance components of single locations (NOR, GBR, CHE, CZE) and multi-location models (ALL). The variation attributed to accession, field heterogeneity as assessed by incomplete blocks (Block), the interaction of accession x location (for multi-location model only) and residual is shown. Traits are crude protein content (CP), date of flowering (DOF), dry matter yield (DMY), plant density in autumn (PD) and plant vigor (VIG). Extensions _Y1 and _Y2 denote the first and second main year of observation, .C1 and .C2 the first and second cut per year, respectively. If the cut number is not specified, average values for VIG and sum of values for DMY over one year were used. Traits that showed low accession variance were accompanied by low heritability. For CP in both cuts, observed heritability values were very low (h^2^<0.25) in all trials, except for the second cut in NOR (Figure 6). Due to the low genetic component of CP, a further detailed analysis of this trait is not provided. Heritability values for DMY showed a very wide range among locations, being lowest in CZE and highest in CHE. Highest heritability values were observed for DOF with values >0.90 except for location CHE (h^2^ = 0.72).

**Figure 6:**
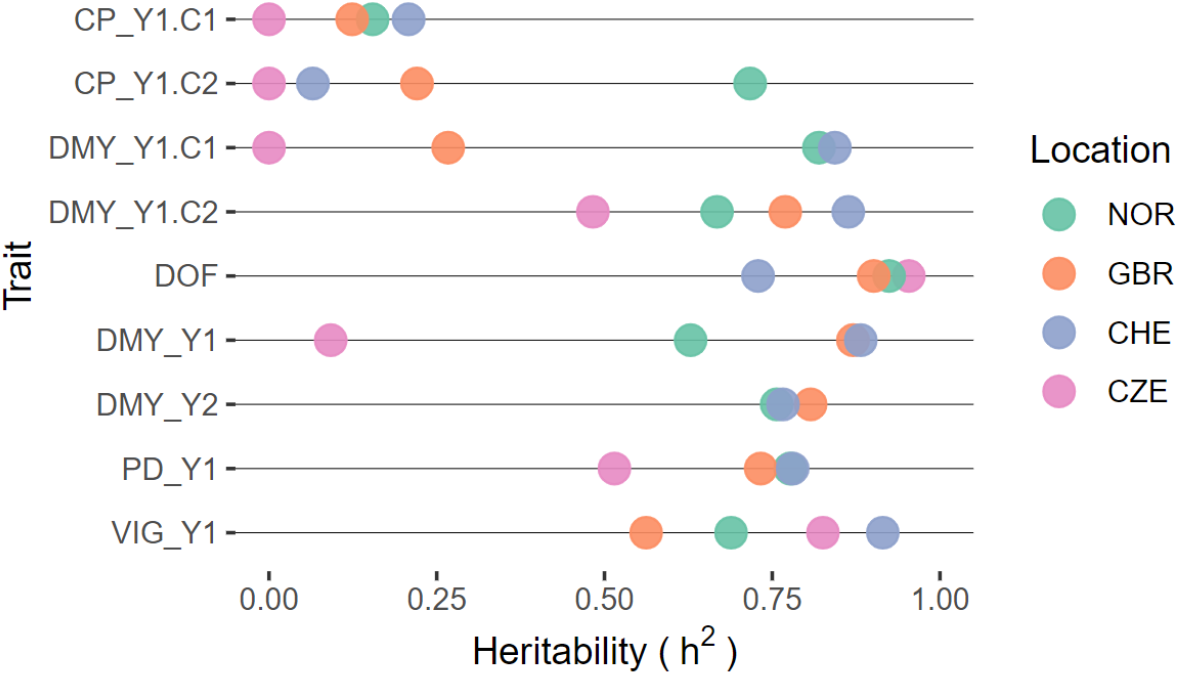
Heritability of selected traits within field trials per location. Traits are crude protein content (CP), dry matter yield (DMY), date of flowering (DOF), plant density in autumn (PD) and plant vigor (VIG). Extensions _Y1 and _Y2 denote the first and second main year of observation, .C1 and .C2 the first and second cut per year, respectively. If the cut number is not specified, average values for VIG and sum of values for DMY over one year were used.

#### 3.2.2 Correlation among locations

The high accession x location interactions observed for different traits (Figure 5), suggest that accessions performed differently at the five locations and that at least one of the trial locations showed a non-positive correlation to the others. Pairwise correlation analyses among locations for traits DMY_Y1, VIG_Y1 and PD_Y1 showed negative correlations of location NOR with all other locations, being significant in some instances (Figure 7). For the same traits, correlation coefficients among the other locations were all positive, being significant in most instances. Thereby, strongest correlations were observed between locations CHE and GBR. For DOF, a trait with only moderate accession x location interaction variance, correlation coefficients were positive among all locations, being significant in most instances (Figure 7).

**Figure 7:**
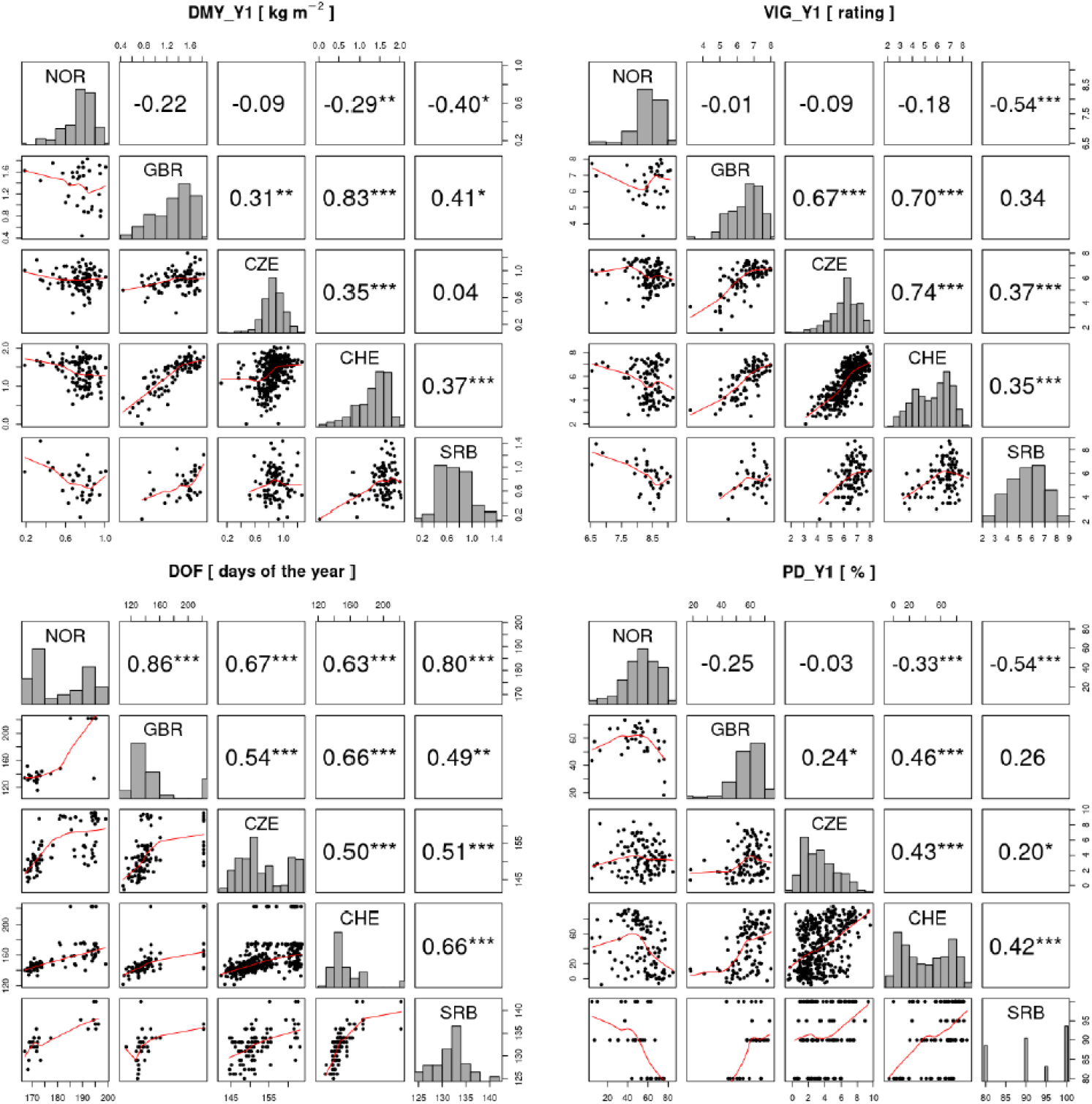
Correlation analysis of the traits total dry matter yield in the first year (DMY_Y1), average vigor in the first year (VIG_Y1), date of flowering (DOF) and plant density after the first year (PD_Y1) in the five field trial locations. In the diagonal, histograms of the trait values for each trial location are shown. Above the diagonal, Pearson correlations between trials are shown and significant correlations are indicated by asterisks (*p<0.05, *** p<0.001). Below the diagonal, trait values of the two trials are plotted against each other with the red line representing the LOESS (locally estimated scatterplot smoothing) line.

#### 3.2.3 PCA of phenotype and disease occurrence

PCA was performed for each trial location using phenotypic data including disease occurrence (Figure 8). Natural disease infections occurred in all trials except location GBR. At location CHE, southern anthracnose was observed at Y0.C2 (h^2^ = 0) and Y1.C4 (h^2^ = 0.40), while powdery mildew was observed at Y0.C1 and Y0.C2 with heritability of 0.75 and 0.80, respectively. In location SRB, brown spot was observed in Y2.C2 (h^2^ = 0.67). In location NOR, clover rot was observed in spring Y1.C1 (h^2^ = 0.59). In location CZE, fusarium plant rot was observed in Y1.C3 (h^2^ = 0.30). Correlations among traits are given in Supplementary Figures S2-6.

**Figure 8:**
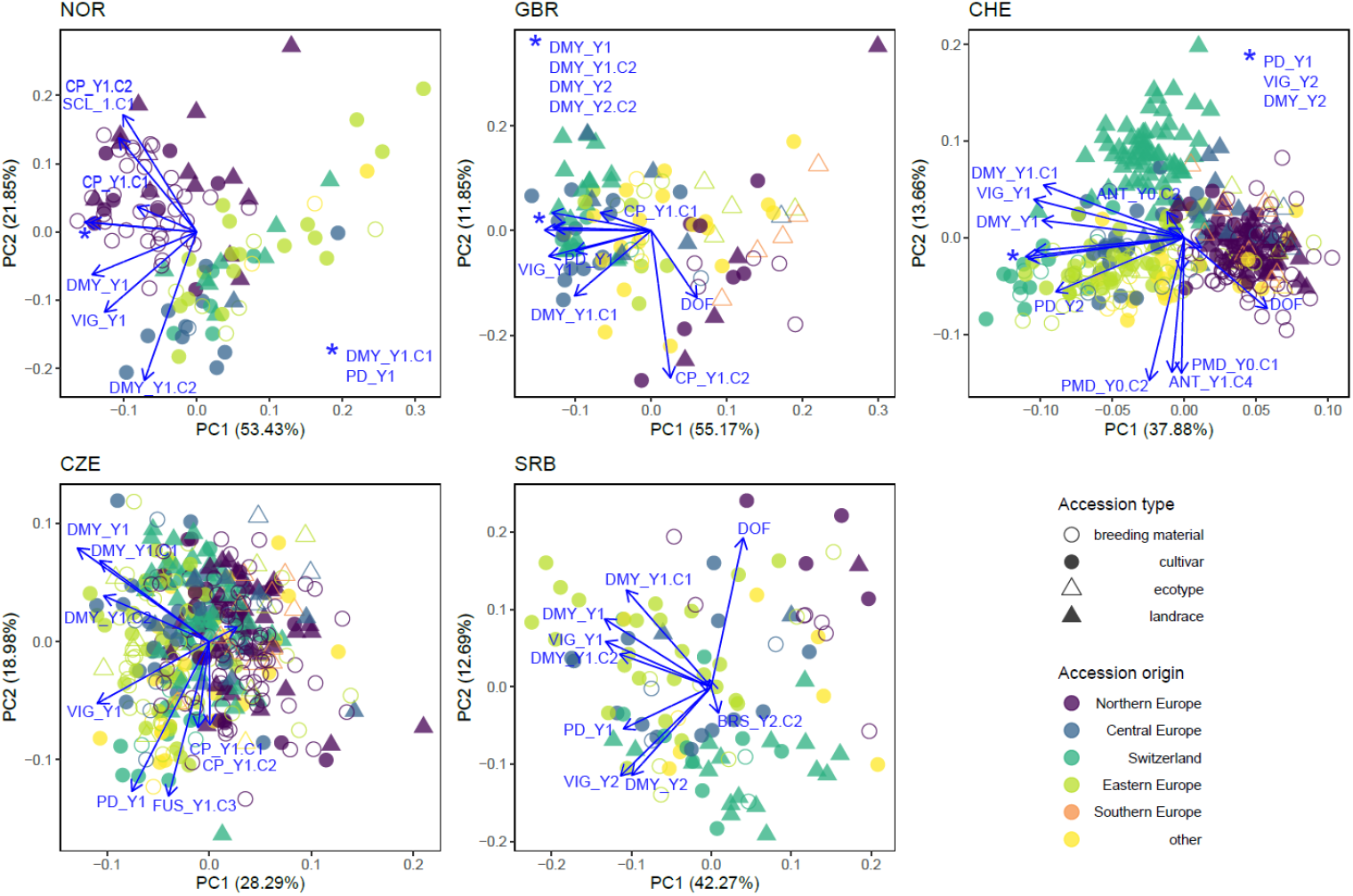
Principal component analysis for each trial location including phenotypic data (DMY_Y1.C1, DMY_Y1.C2, CP_Y1.C1, CP_Y1.C2, DMY_Y1, DMY_Y2, VIG_Y1, VIG_Y2 and PD_Y1), disease scoring (ANT = southern anthracnose, BRS = brown spot, FUS = fusarium plant rot, PDM = powdery mildew, SCL = clover rot) and where they occurred. Colors indicate the region of origin of the accessions and symbols indicate the accession type.

PCA of phenotypic values showed clustering of accessions according to their origin at each trial location. In the NOR trial, the local accessions (accession origin = Northern Europe) clustered in the upper left side and are distinguished from other accessions by an increased resistance to clover rot and an increased CP_Y1.C2. In addition, the local accessions flowered later and expressed high yields mainly in the first, but not in the second cut (high DMY_Y1.C1 and low DMY_Y1.C2). In GBR, PC1 was strongly determined by yield and PC2 by DOF and CP_Y1.C2. Notably, Northern European accessions and ecotypes showed low yield, while Central and Eastern European as well as Swiss accessions were high yielding. In CHE, PC1 mainly distinguished accessions by their sum of DMY per year (DMY_Y1 and DMY_Y2) and yield of the first cut (DMY_Y1.C1), while PC2 represents variation in disease resistance (powdery mildew and southern anthracnose) as well as DOF_Y1. Thereby, Northern European accessions clustered separately from other materials with lower total yields per season, but higher yield in the second cut and later flowering. In CZE, no clear clustering of accessions by origin was observed. Variation in fusarium resistance Y1.C3 and plant density PD_Y1 both correlated with PC2 and are indicative for reduced plant stands after disease infestation. In SRB, the Eastern European accessions are distinguished from the Central European and Swiss accessions mainly by their increased yield and vigor in Y1. A weak trend of Northern European accessions for later flowering and lower vigor in Y1 was observed (Figure 8).

#### 3.2.4 Local adaptation of breeding material and cultivars

To find signs of local adaptation in red clover breeding materials, the performance of accessions belonging to the type ‘breeding material’ or ‘cultivar’ and grouped by their geographic origin, were compared in the different trial locations across Europe (Figure 9). The Northern European accessions flowered later on average in all locations and displayed a clear adaptation to their native northern climate: Northern European accessions showed the highest plant density after Y1 in NOR, while accessions of other origin did not persist very well. However, the Northern European accessions displayed reduced plant densities under non-native conditions and generally did not perform very well concerning yield and vigor. The Central and Eastern European accessions performed comparably well in the trials in CZE, CHE, and SRB for all traits. The Swiss accessions performed slightly better at their origin (CHE) regarding yield and vigor, and much better regarding plant density. However, in the trials in SRB and CZE, the Swiss accessions performed slightly worse compared to the Central and Eastern European accessions regarding yield and vigor. In GBR, the Central European accession performed best, followed by the Swiss and the Eastern European accessions.

**Figure 9.**
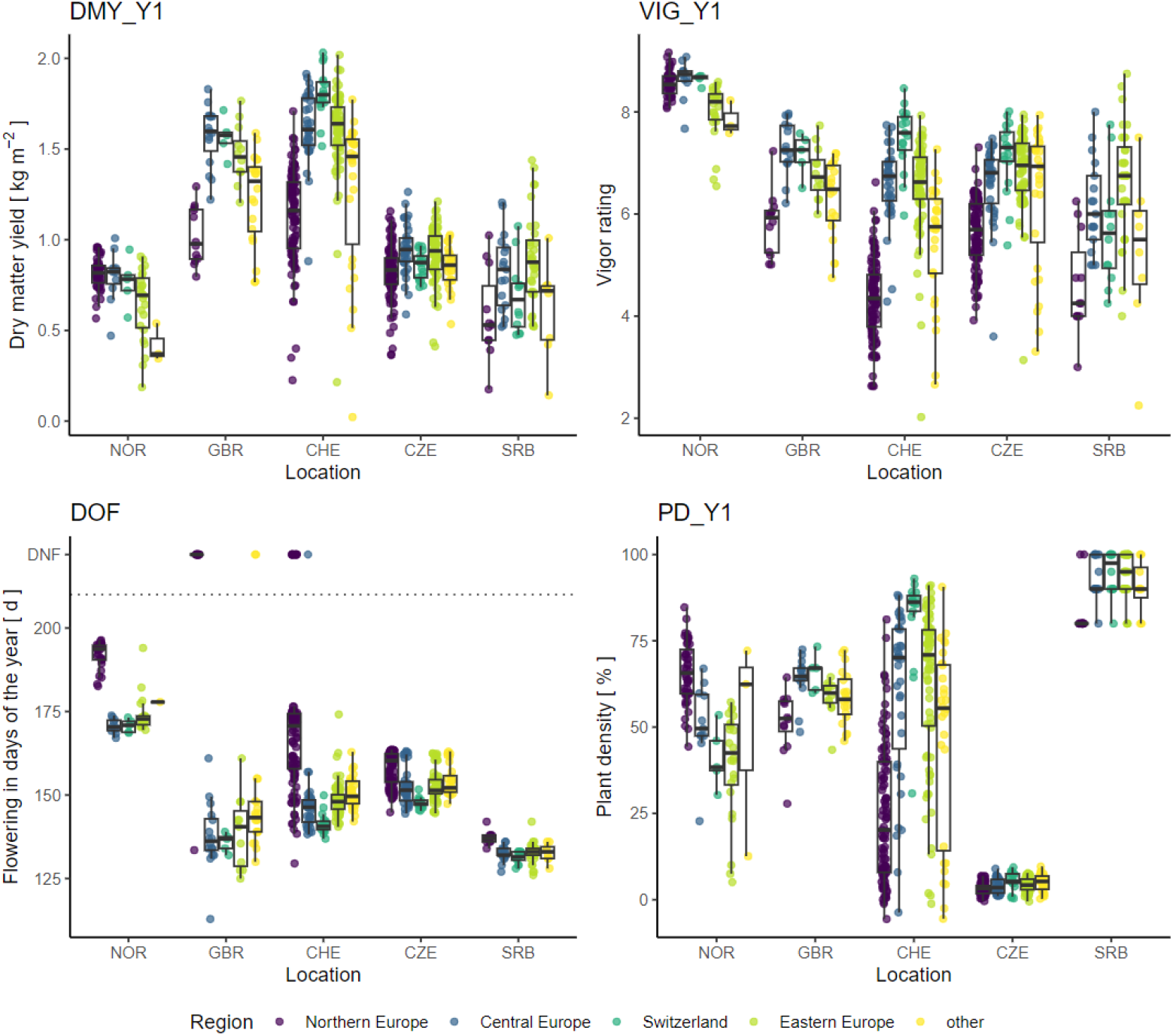
Performance of accessions of the type ‘cultivar’ or ‘breeding material’ grouped by their region of origin. Traits analyzed are total dry matter yield of the first year (DMY_Y1), average vigor of the first year (VIG_Y1), date of flowering (DOF) as well as plant density after the first year (PD_Y1). In the plot of DOF, accessions that did not flower are displayed with a value of 222 and are labelled accordingly (DNF).

#### 3.2.5 Accession x location interaction of accessions tested in all trials

Twenty accessions were tested in all five trials and their accession x location performance was analyzed (Figure 10). The Northern European accessions uniformly showed distinct patterns compared to the remaining groups. Northern European accessions flowered later than the other accessions and, although vigorous and high yielding in NOR, they did not perform well in the other locations. For the accessions from CHE, GBR, Central and Eastern Europe, such a pattern was not observed and within groups, usually good as well as poorly performing accessions were found.

**Figure 10:**
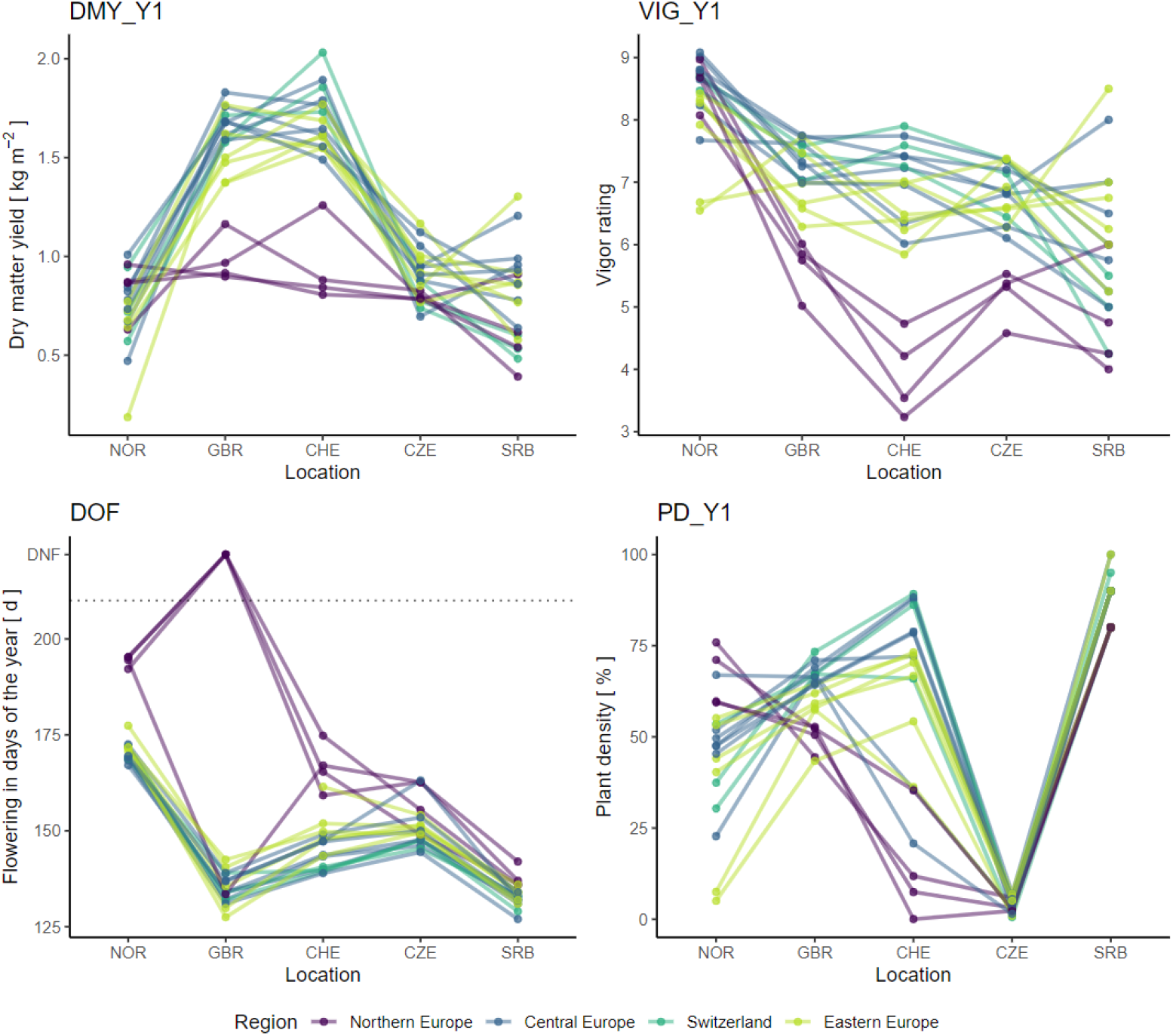
Performance of 20 accessions grown in all five field trials. Traits analyzed are total dry matter yield of the first year (DMY_Y1), average vigor of the first year (VIG_Y1), date of flowering (DOF) as well as plant density after the first year (PD_Y1). In the plot of DOF, accessions that did not flower are displayed with a value of 222 and are labelled accordingly (DNF). Accessions are colored according to their region of origin.

## 4 Discussion

This study, to the best of our knowledge, characterised the most comprehensive collection of red clover accessions to date. By evaluating the collection on field plot level at diverse locations, a wide range of environmental conditions across Europe were covered. Combined with population-level allele frequency genotyping, this allowed to relate genetic diversity to phenotypic performance.

The genetic structure of the EUCLEG panel largely reflected the geographical origin of the accessions. Ecotypes from Southern Europe and some cultivars from non-European countries were clearly separated from the remaining accessions (Figure 2). A similar pattern was observed in an analysis of a global red clover collection, where accessions could be grouped into four regional groups (Asia, Iberia, UK and Central Europe) based on GBS of single plants (Jones et al., 2020). In the present study, accessions from Central and Eastern Europe clustered together, while accessions of Northern Europe and Switzerland each formed distinct clusters. Within the Northern European and Swiss clusters, sub-clusters were identified that distinguish breeding material from landraces (Figure 3). The genetic similarity of landraces and breeding material is in agreement with the historic origin of breeding programs that mostly started with improving local germplasm (Boller et al., 2010). Still, systematic improvement through breeding for traits like disease resistance or yield has led to populations that are distinctly different from the original landraces as indicated by the separate clustering of breeding material and landraces within the Swiss and Nordic accessions (Figure 3). A separation of ecotypes from cultivars has also been demonstrated recently in a study on 29 Nordic red clover accessions using genetic markers (Osterman et al., 2021).

Closer inspection of the genetic structure of the EUCLEG red clover diversity panel reveals a few particularities that may reflect specific breeding histories. For example, the cultivar ‘Merian’ (TP394) from Belgium clustered with the Swiss landraces, indicating the latter being the initial source of breeding germplasm used (Figure 3). According to information from the breeder, the cultivar ‘Merian’ orginates to 75% from ILVO breeding material from before 1970, which may also have included some Swiss landraces (T. Vleugels, personal communication). An interesting cluster of three accessions was observed at the PC1, PC2 coordinates −5, −0.5 (Figure 3): The Swiss landrace ‘Ueberstorf (TP074), as well as the cultivars ‘Grasslands Sensation’ from New Zealand (TP036) and ‘AberClaret’ from the United Kingdom formed a distinct group and the Pearson correlations of their allelic profiles ranged from 0.89 (TP036/TP074) to 0.94 (TP006/TP074; data not shown). ‘Grasslands Sensation’, also known as ‘G40’ or ‘Swiss’ was bred based on four Swiss red clover cultivars, *i.e*. ‘Renova’, ‘Mont-Calme’, ‘Leisi’ and ‘Changins’, all tracing back to Swiss landraces (Kölliker et al., 2003; Claydon et al., 2010). ‘AberClaret’ and the cultivar ‘AberChianti’ (TP007; Figure 3), are advertised as high-yielding up to their fourth and fifth harvest year (Marshall et al., 2017), a typical trait of Swiss ‘Mattenklee’ cultivars and landraces, to which they show considerable similarity in the PCA (Figure 3). The Canadian cultivar ‘Altaswede’ (TP105) clustered within the Northern European accessions. This is in agreement with it being described as ‘single cut’ clover (Valle, 1958), the predominant phenotype of red clover in Northern Europe. Indeed, ‘Altaswede’ was selected by the University of Alberta, Canada, based on Swedish seed stock around 1919 (Aasen and Bjorge, 2009). These examples are clear evidence that red clover breeders sometimes make use of the ‘breeder’s exemption’, *i.e*., the right to use registered cultivars from the market in their own breeding programs and integrate desired traits into their own breeding pools. Nevertheless, the distinct grouping of the accessions according to their region of origin suggests that this is, so far, not frequently done and breeders mostly rely on their own, local genetic resources. This is in contrast with many other breeding programs of crops such as wheat or rice, where the incorporation of foreign material is far more widespread (Garrett et al., 2017; Luttringhaus et al., 2020).

While the characterisation of genetic diversity is an important first step in the efficient utilization of genetic resources, targeted incorporation of germplasm into the breeding process requires detailed phenotypic characterisation. Conducting field trials with perennial forage legumes is challenging. First, the establishment phase is critical to ensure reliable data in the following years. Second, pest, disease or weed occurrence can substantially reduce the plant density after each cut or during harsh winters (Boller et al., 1998; Öhberg et al., 2008; Zanotto et al., 2021), having a strong impact on the data assessed in subsequent cuts and growing seasons. Although the multi-site trials in this study were all managed by experienced red clover researchers and field teams, there were some setbacks during the trial period. The trial in CZE had to be abandoned after year one, as the damage caused by mice and *Fusarium* spp. was too severe to get reliable data in subsequent years. Several other trials experienced disease pressures that resulted in the loss of several accessions and missing data in the following cuts. Despite the difficult conditions encountered at some trial locations, heritability values of field trials were, with a few exceptions, at a high level (h^2^ >0.6, Figure 6) and similar to heritability values reported in other perennial forage crops (Annicchiarico et al., 1999; Han et al., 2006; Li et al., 2015).

A trait showing low heritability across all trial locations was CP. The reason for this is not entirely clear, but may be associated with the low overall variability for this trait (coefficient of genetic variation (CVg) = 0.07-0.18, Supplementary Table S2). Only the heritability of CP of the second cut (CP_Y1.C2) for trial site NOR was satisfactory with h^2^ = 0.72. It was highly correlated with DOF (r = 0.76, Supplementary Figure S2) and negatively correlated to DMY_Y1.C2 (r = −0.28). Hence, accessions that had a high CP in C2 generally flowered later, yielded more in C1 and less in C2 and may have concentrated protein levels in C2 (data not shown). Despite several publications reporting red clover forage quality values (Schubiger et al., 1998; Hoekstra et al., 2018), to our knowledge no heritability values have been reported so far. Thus, it remains unclear whether the heritability is truly low or, whether there was not enough variation in the accessions studied. The latter has previously been reported for alfalfa, where limited variability led to low narrow sense heritability of CP (Guines et al., 2002). In addition, the CP levels of red clover are already high compared to forage grasses (Schubiger et al., 1998) and therefore not a primary breeding goal. In alfalfa, leaf-stem ratio influences forage quality and is an important breeding aim (Annicchiarico, 2015).

During the experiment, the plant density of several red clover accessions declined at specific locations. In NOR, the plant density in autumn of Y1 was on average 39% for Eastern European accessions compared to 63% for native Nordic accessions (data not shown). In comparison, in SRB the plant density was on average 83% for Nordic accessions compared to over 90% for accessions from other regions. The overall high plant density in SRB after Y1 (Figure 9) is likely caused by the late sowing (autumn compared to spring for other trials) and the mild winter following sowing. In the Swiss trial, which experienced high southern anthracnose disease pressure, average plant densities after Y1 were highest for local, American and Eastern European accessions (62%, 57% and 59%, respectively) and lowest for Northern European accessions (21%), which is in agreement with the level of southern anthracnose resistance observed for these accessions in artificial inoculation experiments (Frey et al., 2022). The low plant density after Y1 resulted in non-satisfactory data quality in the second year for many accessions. Often, accessions that failed to persist were of foreign origin and not adapted to the conditions of the trial location, which indicates a strong effect of geographical origin on performance. Indeed, the clustering by geographical origin of accessions was also apparent when phenotypic data per trial was analysed (Figure 8). The genetic distinctness of these populations is, thus, manifested in their phenotypic performance. It also indicates a degree of local adaptation among accessions originating from similar environments.

In 1937, Pieters and Hollowell reported that “claims of superiority are made for all of these regional strains, and the evidence of comparative trials shows that in most cases each such regional variety is superior to others in the environment where it was developed” (Pieters and Hollowell, 1937). The present work provides strong support for this hypothesis with trial locations covering a wide range of eco-environmental conditions relevant for European red clover cultivation areas. Breeding material and cultivars from Northern Europe showed best performance at the NOR location with regard to DMY and vigor (Figure 9). This trend of local adaptation with accessions performing well at the location where they were bred was also observed for other trial locations and is in agreement with previous findings (Valle, 1958; Rosso and Pagano, 2005). In the CHE trial, Swiss breeding materials and cultivars showed best performance with regard to DMY_Y1 and VIG_Y1, while in the GBR trial Central European accessions performed best. In CZE and SRB, Eastern European breeding materials performed best (Figure 9). As highlighted above, performance of red clover is often driven by persistence, and the plant density of poorly adapted accessions declines throughout the growing season and/or winter. While red clover has a natural senescence, its persistence can be improved by improving resistance for relevant diseases that occur in the growing location (Taylor, 2008). Depending on the region, different traits are relevant for persistence and hence have resulted in accessions adapted to the location of origin.

The five trial locations cover a wide range of latitudes and climate zones, and large accession x location interactions were observed. From correlation analysis among locations (Figure 7), we could show that especially the Northern European trial location (NOR) differed from the remaining sites. Negative correlation coefficients indicated the presence of cross-over interactions, *i.e*. that ranking of accessions is reversed in NOR compared to other locations. This observation clearly highlights the need for locally adapted breeding programs. We observed a strong adaptation of Northern European accessions to the prevailing conditions and/or management. According to Boller et al. (2010), European cultivars of red clover can be grouped by flowering time: early-flowering types adapted to southern latitudes are capable of rapid regrowth and continuous flowering, while late-flowering types adapted to northern latitudes remain vegetative after the first cut. This may explain the poor performance of Northern European accessions of red clover in the more southern locations, where the cutting regime is optimized for fast regrowing types. At these locations, the first cut is most likely too early for Northern European accessions, which show lower yield as they are not able to resume generative growth afterwards. In their native habitat of NOR, the first cut happened 1-1.5 months later than in the remaining trial locations and allowed the Northern European accessions to reach their full yield potential.

Due to the natural expansion across Eurasia and Northern Africa and the recent domestication at several locations (Kjærgaard, 2003), red clover has evolved as a very diverse crop. The outcrossing nature of red clover has, in addition to the large between-accession diversity, resulted in an even higher within-accession diversity (Kölliker et al., 2003; Dias et al., 2008; Pagnotta et al., 2010; Collins et al., 2012). This allows red clover accessions to rapidly adapt to new challenges. Being a perennial species, adaptation in red clover has even been detected during continuous cultivation without allowing reproduction. Collins et al. (2012) found considerable genetic shifts within accessions grown at locations different than their origin after three to five years of continuous cultivation. Especially the Swedish environment was very selective and composite accessions became more similar to the Swedish accessions than to their original accession (Collins et al., 2012).

As indicated by the large accession x location interactions observed in our study, breeding a ‘one-fits-all’ cultivar seems to be very unrealistic, due to the very variable environmental conditions demanding different adaptive traits from the genetic material as discussed above. However, there are trial locations that were more similar to each other based on assessed trait values and cultivars being able to perform well in such regions are more likely to be found. Growing a reduced set of diverse accessions at more locations throughout Europe might help to define such macro-environments representing the different red clover growing conditions. Knowledge of available genetic resources and their characteristics is key for breeding red clover well adapted to environmental conditions. This will become even more important in view of changing climatic conditions. For example, with increasing temperatures diseases adapted to warmer climates like southern anthracnose, which has already extended its local distribution (Jacob et al., 2016; Hartmann et al., 2022), will most likely continue to spread to northern latitudes. Materials resistant to this disease might be a valuable source to complement the genetics from other regions not yet affected. The present work will serve as a valuable basis to identify interesting materials expressing desired characteristics. In conclusion, the detailed description of the EUCLEG red clover diversity panel provides an important source for further targeted improvement of red clover. The knowledge gained about the importance of local adaptation and genetic differentiation will help to design novel breeding strategies that consider broader scale adaptation of future cultivars.

## 5 Supplementary Material

Supplementary Table S1: Accessions, origins and experimental sets

Supplementary Table S2: Trait values and their statistical distribution at the five field locations

Supplementary Figure S1: K-means clustering of principal components

Supplementary Figure S2-6: Correlation of traits within locations

## 6 Data Availability Statement

The data presented in this study are openly available at zenodo.org (https://doi.org/10.5281/zenodo.7447473)

## 8 Author Contributions

Study design and curation of the accession collection: CG, RK, AP; Generation of phenotypic data: CG, HA, JR, LJ, LS; Generation of genotypic data: LS, TR, LAF; Data analysis and interpretation: MMN, RK, CG, LAF; Drafting of the manuscript: MMN, RK, CG; All authors contributed to writing and read and approved the final version of the manuscript.

## 10 Acknowledgments

We thank the EUCLEG consortium for the valuable training and exchange sessions and all institutions providing red clover seeds, including: AgResearch (NZL), Agricultural Res. Ltd. (CZE), Agroscope (CHE), Boreal (FIN), DLF Seeds (CZE), DSV (GERDE), Graminor (NOR), HBLFA (AUT), Hokkaido Ag. Res. (JPN), IBERS (GBRUK), IFVCNS (SRBRS), ILVO (BEL), INRA (FRA), Lantmännen (SWE), NordGen (SWE), PGG Wrightson (NZL), RAGT2n (FRA), and USDA (USA). In addition, we would like to thank the technical staff at Aberystwyth University, Agroscope, DLF Seeds, Graminor, and Institute for Forage Crops Kruševac for their management of the field trials.

