## Supplementary Figures for "Multi-location trials and population-based genotyping reveal high diversity and adaptation to breeding environments in a large collection of red clover"

### *Supplementary Material*

#### 1 Supplementary Figures

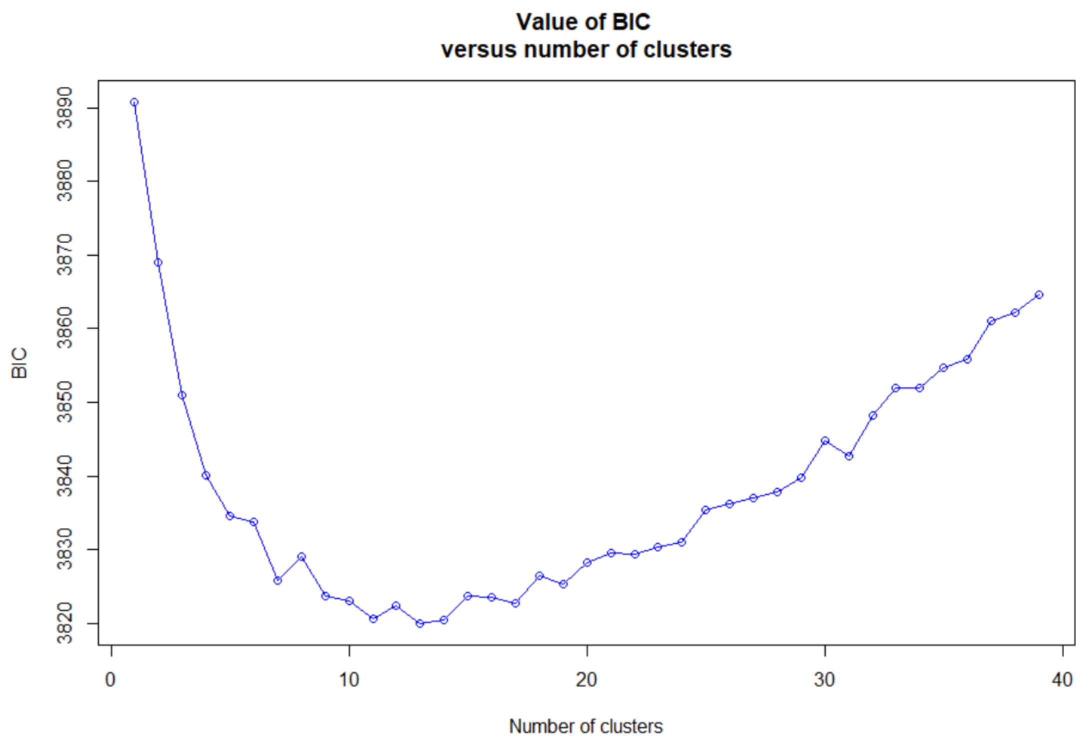


**Supplementary Figure S1.** Successive K-means clustering of principal components derived from 392 red clover accessions and 20,137 SNP markers.

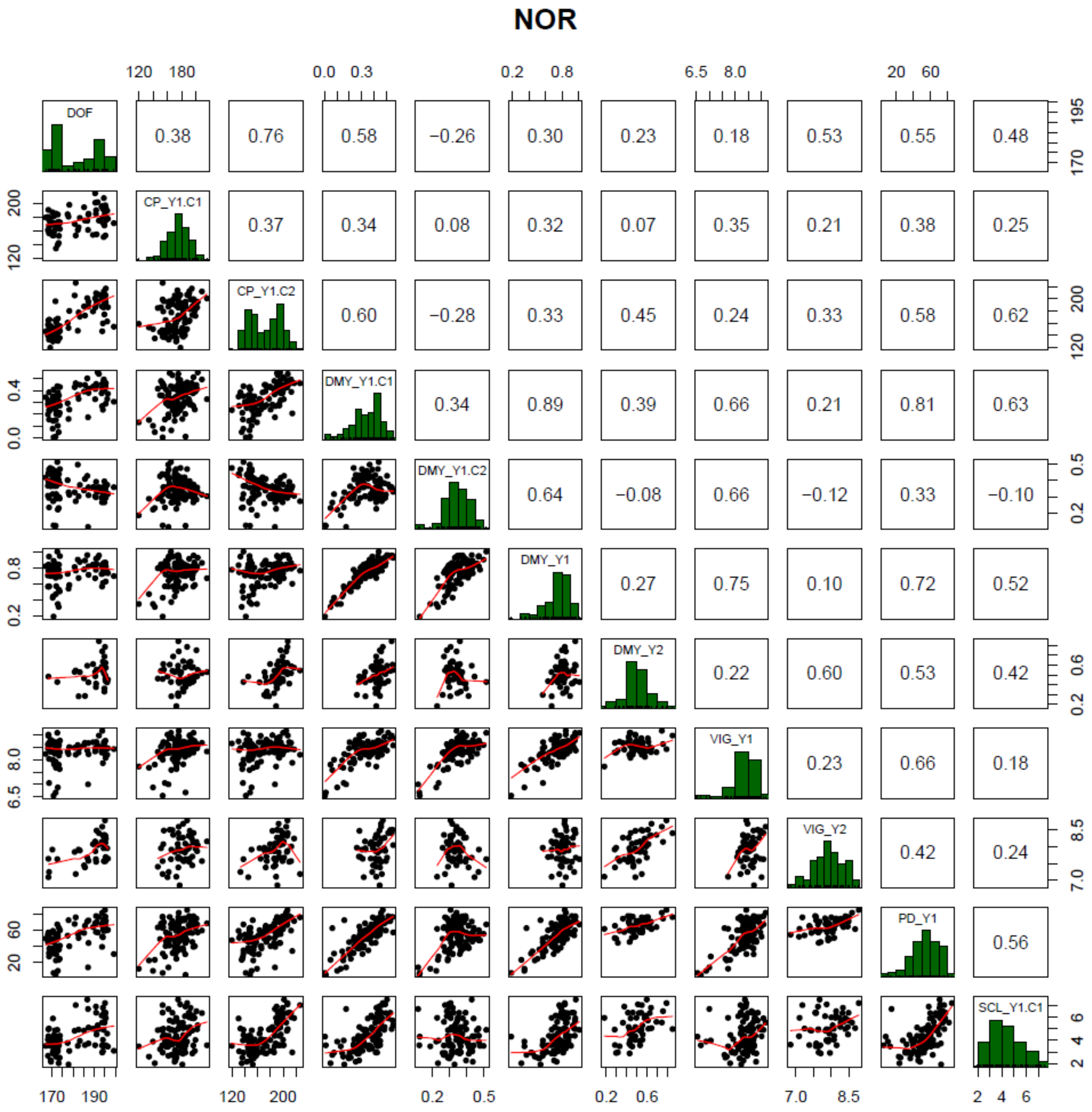


**Supplementary Figure S2.** Correlation of traits measured in the field trial in Bjørke, Norway (NOR). Traits were days to flowering (DOF), crude protein content (CP), dry matter yield (DMY), vigor (VIG), plant density in autumn (PD) and sclerotinia root rot (SCL). Extensions _Y1 and _Y2 denote the first and second main year of observation, .C1 and .C2 the first and second cut per year, respectively. If the number of cut is not specified, average values for VIG and sum of values for DMY over one year were used.

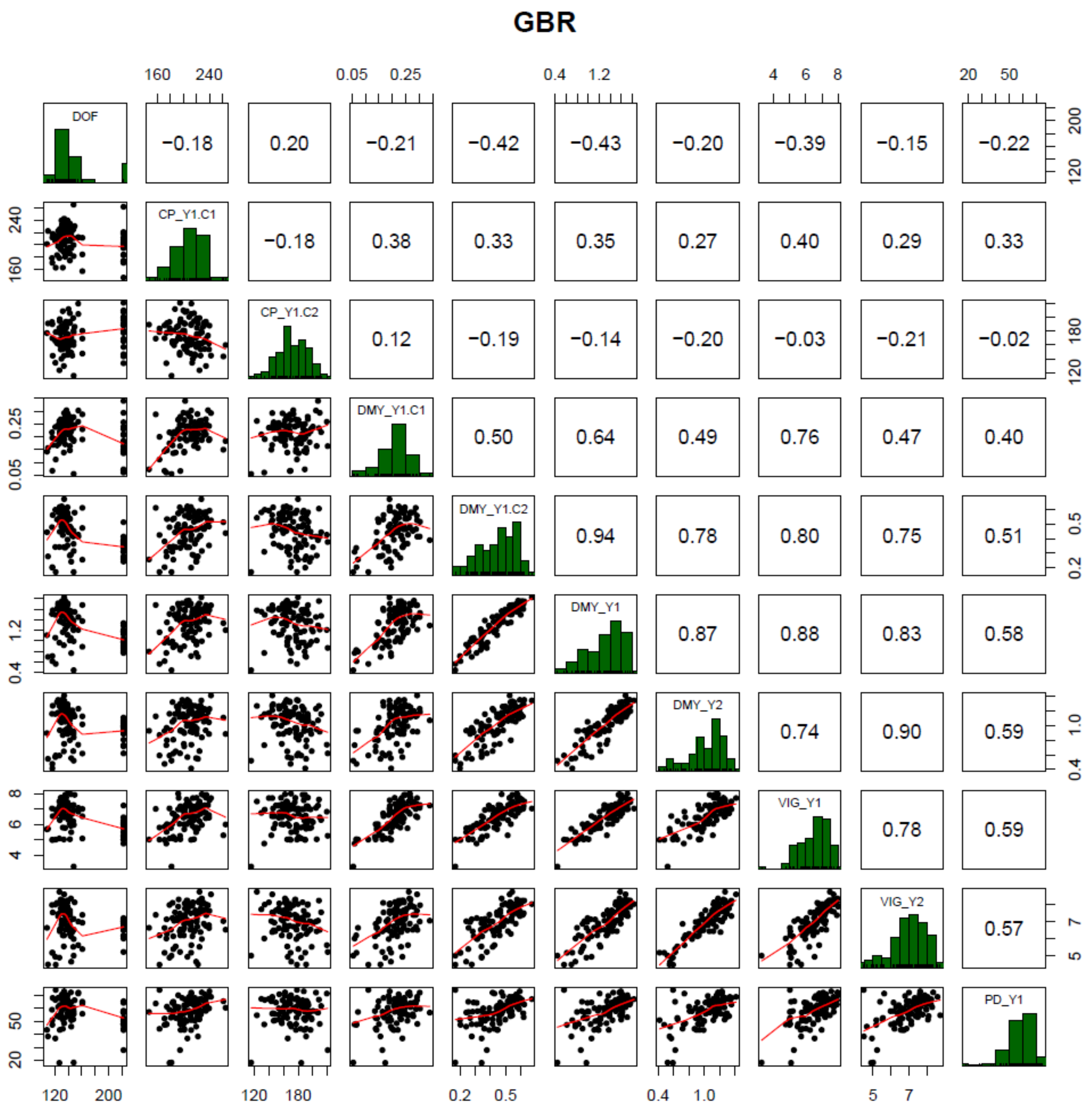


**Supplementary Figure S3.** Correlation of traits measured in the field trial in Aberystwyth, Wales (GBR). Traits were days to flowering (DOF), crude protein content (CP), dry matter yield (DMY), vigor (VIG) and plant density in autumn (PD). Extensions _Y1 and _Y2 denote the first and second main year of observation, .C1 and .C2 the first and second cut per year, respectively. If the number of cut is not specified, average values for VIG and sum of values for DMY over one year were used.

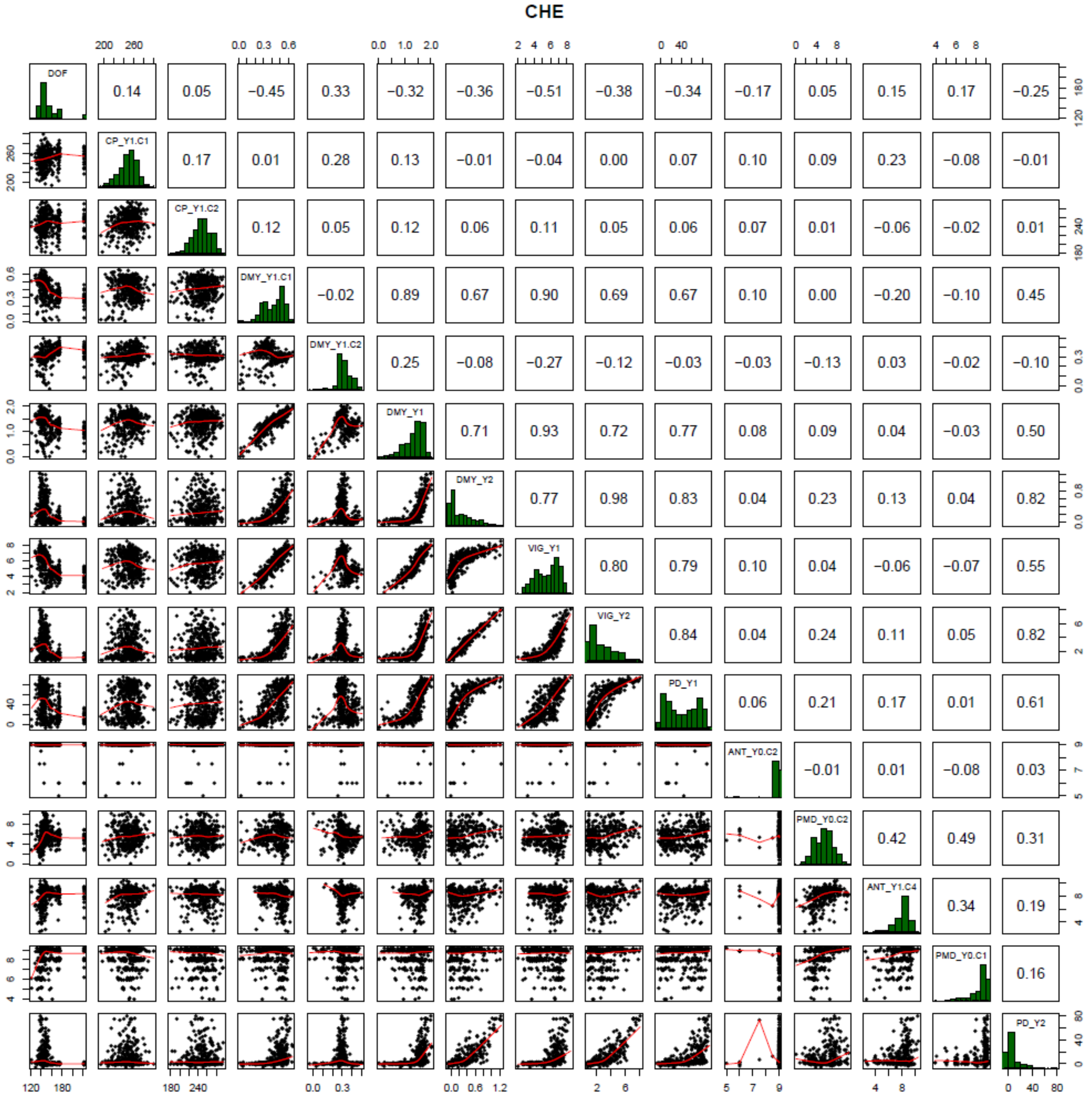


**Supplementary Figure S4.** Correlation of traits measured in the field trial in Tänikon, Switzerland (CHE). Traits were days to flowering (DOF), crude protein content (CP), dry matter yield (DMY), vigor (VIG), plant density in autumn (PD), anthracnose (ANT) and powdery mildew (PMD). Extensions Y0, Y1 and Y2 denote the year of sowing, first and second main year of observation, .C1, C2 and .C4 the first, second and fourth cut per year, respectively. If the number of cut is not specified, average values for VIG and sum of values for DMY over one year were used.

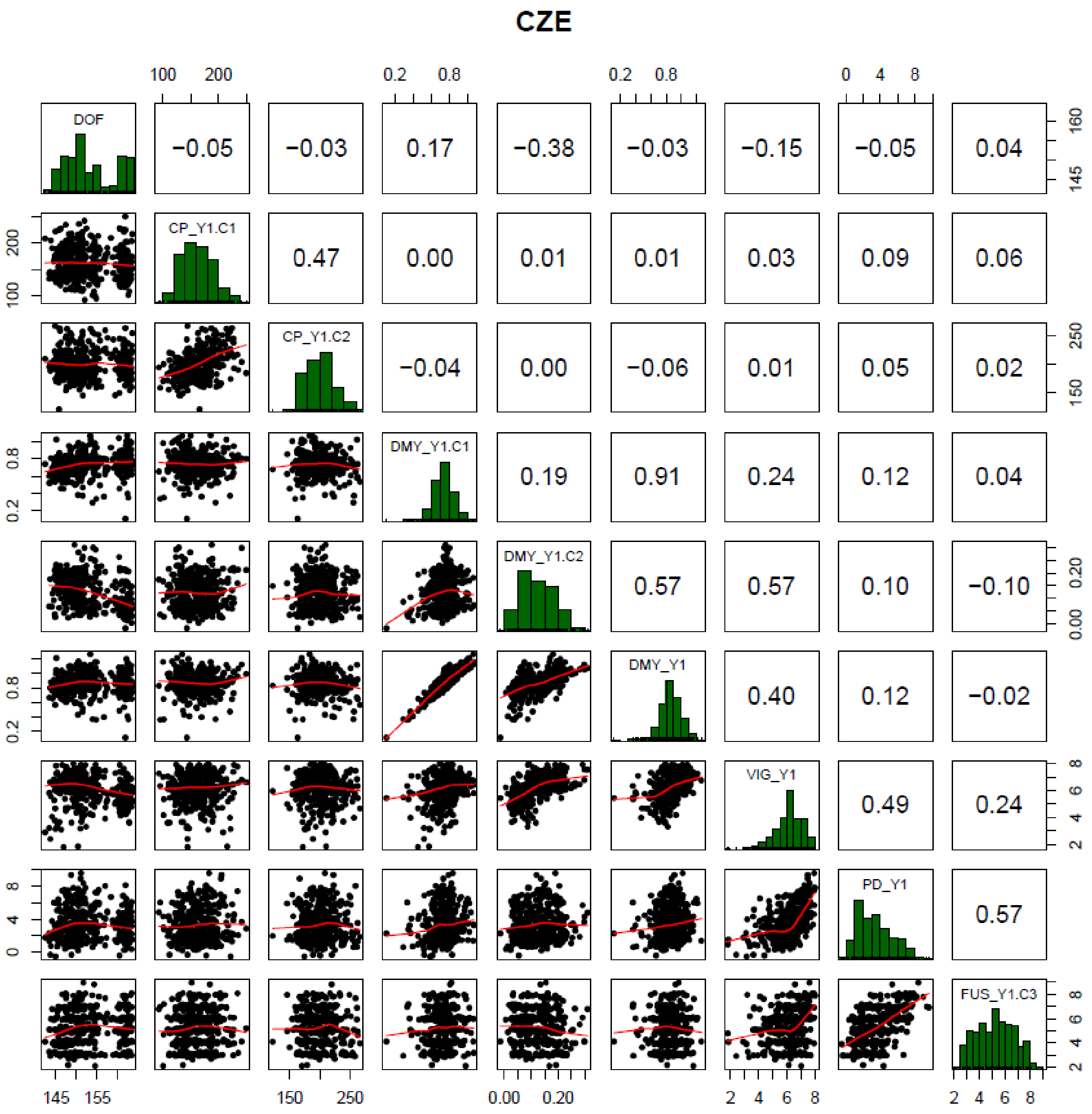


**Supplementary Figure S5.** Correlation of traits measured in the field trial in Hladké Životice, Czech Republic (CZE). Traits were days to flowering (DOF), crude protein content (CP), dry matter yield (DMY), vigor (VIG), plant density in autumn (PD) and fusarium plant rot (FUS). Extensions _Y1 and _Y2 denote the first and second main year of observation, .C1, C2 and .C3 the first, second and third cut per year, respectively. If the number of cut is not specified, average values for VIG and sum of values for DMY over one year were used.

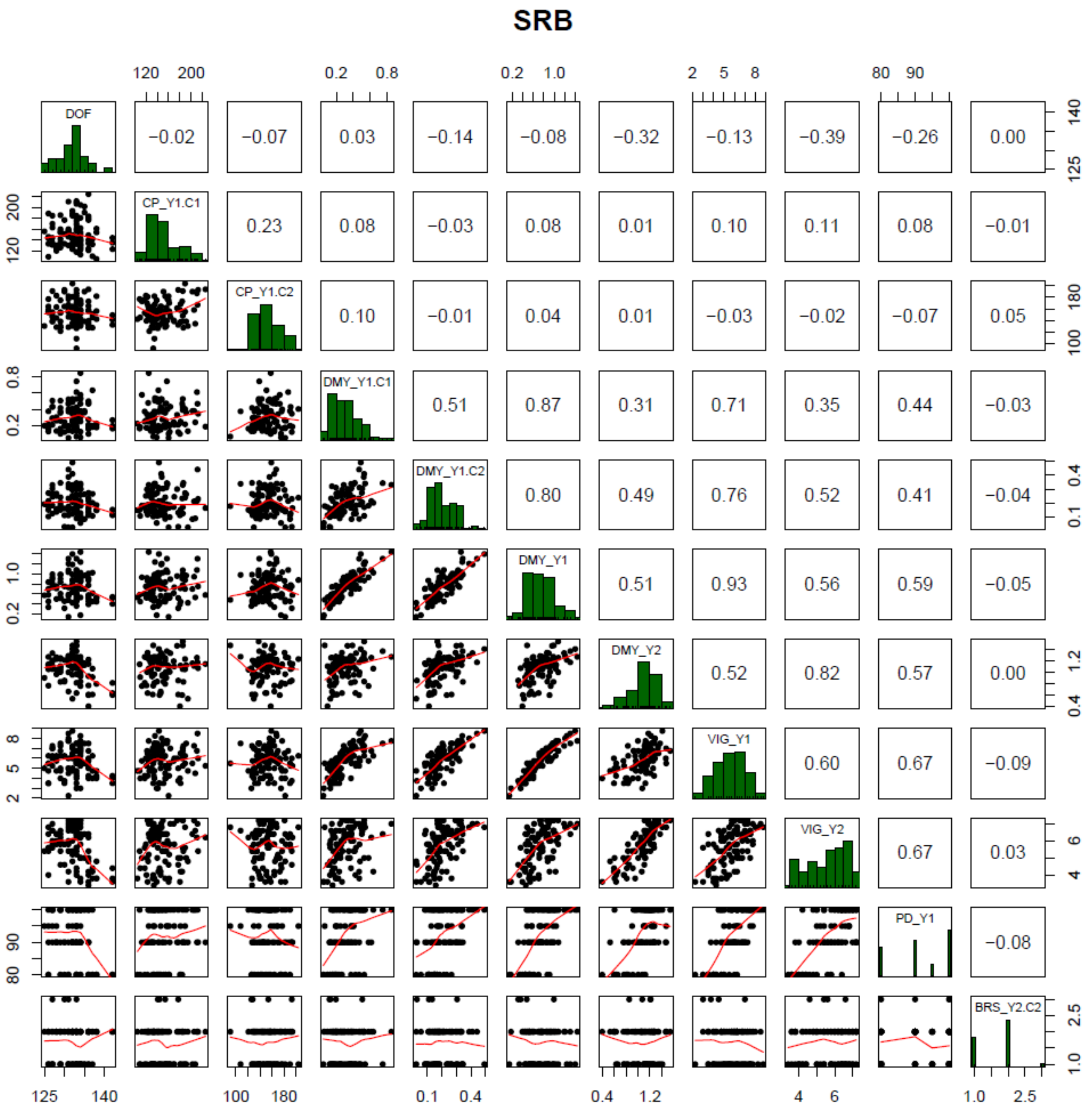


**Supplementary Figure S6.** Correlation of traits measured in the field trial in Kruševac, Serbia (SRB). Traits were days to flowering (DOF), crude protein content (CP), dry matter yield (DMY), vigor (VIG), plant density in autumn (PD) and brown spot (BRS). Extensions _Y1 and _Y2 denote the first and second main year of observation, .C1 and .C2 the first and second cut per year, respectively. If the number of cut is not specified, average values for VIG and sum of values for DMY over one year were used.
